# Vitamin D3 attenuates nitrogen mustard-induced dermal toxicity by enhancing microbial butyrate production via the intestinal VDR-α-defensin signaling pathway

**DOI:** 10.64898/2026.03.24.713897

**Authors:** Xunhu Dong, Ying He, Xiaofeng Hu, Zhe Zhang, Feng Ye, Hongying Chen, Min Qin, Xiaogang Wang, Yuanpeng Zhao, Guorong Dan, Jiqing Zhao, He Tang, Yan Sai, Aiping Wang, Hongbing Song, Zhongmin Zou, Mingliang Chen

**Author notes:** Correspondence: Aiping, Wang M.D., Ph.D., Professor, Department of Frigid Zone Medicine, College of High Altitude Military Medicine, Army Medical University, 30 Gaotanyan Street, Shapingba District, Chongqing 400038, China. Hongbin Song, Ph.D., Professor, Chinese PLA Center for Disease Control and Prevention, No.20 Dongdajie Street, Fengtai District, Beijing 100071, China. Zhongmin Zou, M.D., Ph.D., Professor, Institute of Toxicology, School of Military Preventive Medicine, Army Medical University 30 Gaotanyan Street, Shapingba District, Chongqing 400038, China. Mingliang Chen, M.D., Ph.D., Associate professor, Institute of Toxicology, School of Military Preventive Medicine, Army Medical University, 30 Gaotanyan Street, Shapingba District, Chongqing 400038, China. These authors contribute equally to this work (Xunhu Dong, Ying He, Xiaofeng Hu, Zhe Zhang, Feng Ye).

## Abstract

Nitrogen mustard (NM)-caused severe cutaneous damage lacks effective targeted therapies. Vitamin D3 (VD3) shows promise as a therapy for NM-induced dermal toxicity; however, the underlying mechanisms remain elusive. Herein, we initially confirmed that NM induced gut flora dysbiosis, characterized by a decrease of *Akkermansia muciniphila (AKK)* abundance, thereby leading to butyrate reduction. Antibiotics (ABX) significantly promoted NM-induced skin injury, whereas fecal microbiota transplantation of the controls’ feces (HC-FMT) or *AKK* administration attenuated NM-induced dermal toxicity. HC-FMT or *AKK* significantly increased butyrate levels in feces and serum of NM-treated mice. Butyrate notably attenuated ABX-caused acceleration of NM-induced skin injury. Meanwhile, NM markedly decreased the expression of α-defensins, MMP7, and VDR. NM failed to further decrease *AKK* abundance and BA contents in intestinal MMP7-deficient mice, which was abolished by human alpha defensin 5 (HD5) overexpression. And intestinal MMP7 deficiency enhanced NM-caused skin injury, which was markedly attenuated by HD5 overexpression, *AKK* transplantation, or BA supplementation. Moreover, NM also failed to further reduce MMP7 and α-defensin expression, *AKK* abundance, and butyrate levels in intestinal VDR-silenced mice. Finally, VD3 remodeled the gut microbiome particularly enriching *AKK,* increased butyrate contents and promoted the expression of α-defensins, MMP7, and VDR, thereby attenuating NM-induced skin damage. The protective effect of VD3 against NM-caused dermal toxicity was abolished by either ABX or intestinal-specific knockdown of MMP7 or VDR in mice; however, this impairment was reversed by butyrate or *AKK*. In conclusion, VD3 attenuated NM-caused dermal toxicity by promoting BA production via remodeling the gut microbiota, and this effect was partially mediated by the intestinal VDR-α-defensin signaling pathway. These highlight that targeting the gut flora or supplementing with BA could be potential therapies for NM-induced dermal toxicity.

## 1. Introduction

Nitrogen mustard (NM), a classic vesicant agent, has been shown to exert adverse effects on multiple organ systems. In particular, the skin represents the principal and most sensitive target for NM.^1,2^ NM-induced skin injury manifests as erythema, blistering, ulceration, and eschar formation and is characterized by prolonged healing, susceptibility to necrosis and infection.^1,2^ At present, effective treatments are unavailable due to an incomplete mechanistic understanding.

The gastrointestinal microbiota refers to microorganisms that inhabit the gastrointestinal tract with diverse properties and functions.^3^ Recently, the gut microbiota has been shown to play important roles in the pathogenesis and treatment of various skin disorders and injuries.^4^ The gut microbiota communities of individuals with atopic dermatitis,^5^ psoriasis,^6,7^ acne vulgaris,^8^ diabetic foot ulcers,^9^ and photoaging^10^ are different from those of the controls, and certain bacterial taxa have been identified as pivotal constituents of the microbiome that modulate cutaneous disorders and wound healing through orchestrating metabolic, epigenetic, and immune pathways.^11,12^ Ablation of specific subsets of core gut bacteria and/or supplementation with probiotics inhibit the pathogenesis of cutaneous lesions and trauma.^13,14^ Accordingly, the microbiota-targeted therapies are considered as potentially effective strategies for dermatoses and cutaneous injuries. Noteworthy, researchers have shown that germ-free mice confer resistance to NM-induced delayed lethality, suggesting that the gut microbiota plays a critical role in the systemic toxicity of NM.^15^ However, the contribution of the gut microbiota, especially core microbial taxa, to NM-induced dermal toxicity and its mechanistic basis remains poorly understood.

Accumulating evidence demonstrates that bacterial metabolites are crucial mediators for the gut microbiota’s effects on skin disorders via the gut-skin axis.^11^ Among which, short chain fatty acids (SCFAs), including acetate, propionate, and butyrate (BA) produced by microbial fermentation of dietary fiber, are considered the most important.^16,17^ Reportedly, the gut microbiota-derived SCFAs can access the circulation and modulate skin cells, helping to regulate the skin barrier and inflammatory responses.^4^ Reduced fecal and serum SCFA concentrations are characteristic of atopic dermatitis, whereas a diet rich in fermentable fiber mitigates systemic allergen sensitization and disease severity by promoting SCFA (especially BA) generation.^18,19^ Acetate and propionate levels are decreased in patients with psoriasis, and SCFA treatment significantly attenuates skin thickening and IL-17 levels in an imiquimod-induced murine model of psoriasis.^20,21^ Moreover, Zhou et al. recently confirmed that specific Traditional Chinese Medicine improves diabetic wound healing through elevating microbial SCFA concentrations.^22^ Meanwhile, melatonin ameliorates sleep deprivation-induced skin injury by modulating the gut microbiota to generate propionate.^23^ More recently, the combination of galacto-oligosaccharide with collagen-tripeptide or *Bifidobacterium longum* has been found to protect the skin against ultraviolet B (UVB)-induced photoaging through increasing SCFA contents.^24,25^ These results indicate that the intestinal flora participates in the pathogenesis of gut-related skin disorders and injuries primarily through regulating SCFA production. However, the exact role of microbial SCFAs in NM-caused dermal toxicity needs further studies.

α-Defensins, predominantly produced by Paneth cells in the small intestine (SI), have been recognized as key regulators of the gut microbiota by participating in pathogen clearance and symbiotic balance of the intestinal environment.^26,27^ Reduced or dysfunctional α-defensins have been implicated in various dysbiosis-related disorders including Crohn’s disease,^28^ obesity,^29^ depression^30^ and alcoholic steatohepatitis.^31^ Sakura et al. also found that neonicotinoid pesticide clothianidin exposure inhibits α-defensin generation thereby inducing gut microbiota dysbiosis in mice.^32^ Wu and his colleagues identified that α-defensins attenuate radiation-caused intestinal injury via remodeling the gut microbiota.^33^ These establish α-defensins as key regulators in health and disease, especially physical or chemical stimuli-induced injury, through modulation of the intestinal flora. Moreover, vitamin D receptor (VDR), highly expressed in SI tissue, serves as a key regulator of α-defensin generation.^34^ VDR knockout decreases the expression of α-defensins and matrix metalloproteinase 7 (MMP7, centrally involved in α-defensin maturation) in the ileum, resulting in intestinal flora dysbiosis that is reversed by synthetic α-defensin 5 supplementation in mice.^35,36^ Meanwhile, *Lactobacillus plantarum* induces α-defensin secretion in a VDR-dependent manner, thereby attenuating *Salmonella*-induced colitis.^37^ These data reveal that VDR plays a key role in α-defensin production and subsequent gut microbiota homeostasis.

Vitamin D3 (VD3) is a skin-synthesized steroid hormone that exerts its actions via VDR interaction.^38^ Growing evidence shows that VD3 is a viable therapeutic intervention for NM-induced skin injury, yet the underlying mechanisms remain unclear. Single-dose intraperitoneal injection of VD3 has been found to attenuate NM-caused dermal toxicity by suppressing NLRP3 inflammasome activation^39^ and ferroptosis^40^ in keratinocytes. Moreover, many human studies have confirmed that VD supplementation significantly alters microbiome composition.^41^ Recently, Xiang et al. reported that VD3 improves high-fat diet-caused obesity parameters through reshaping the gut microbiota in mice.^42^ Meanwhile, VD3 restores lipopolysaccharide (LPS)-induced gut flora dysbiosis thereby alleviating inflammation in the colon epithelium.^43^ It has also been found that VD3 functions through epithelial VDR to alter the gut microbiota, thereby permitting tumor control.^44^ These published data implicate the gut microbiota in VD3’s beneficial effects. Nevertheless, the specific role of the gut microbiota in VD3-mediated protection against NM-caused dermal toxicity requires further investigation.

Our results demonstrated, for the first time, that VD3 attenuated NM-caused dermal toxicity by promoting BA production via modulating the gut microbiota especially enriching *A. muciniphila*, and this effect was partially mediated by the intestinal VDR-α-defensin signaling pathway. Our findings are important complements to previous studies regarding the role of the gut microbiota in skin injuries and provide a new molecular mechanism of VD3 that might be utilized for NM-induced skin injury treatment. These highlight that targeting the gut flora or supplementing with BA could be potential therapies for NM-caused dermal toxicity.

## 2. Materials and methods

### 2.1 Mice

Six-week-old male and female C57BL/6J mice and MMP7^−/−^ mice were purchased from The Jackson Laboratory (USA). Transgenic mice expressing human defensin 5 (HD5) were designed and generated by Vital River Laboratory (Beijing, China). These mice were subsequently crossed with MMP7^−/−^ mice to generate MMP7^−/−^;Tg(HD5) mice. All mice were maintained under specific pathogen-free conditions on a strict 12-hour reverse light-dark cycle, with free access to sterile water and a commercial standard diet. All animal experiments included age-matched cohorts of males and females and were approved by the Laboratory Animal Welfare and Ethics Committee of the Army Medical University.

### 2.2 NM-induced dermal injury model

Mice were anesthetized by intraperitoneal injection of pentobarbital sodium (50 mg/kg), dorsal hair was removed using electric clippers followed by a depilatory cream. After 48 hours, the dorsal skin of mice (n =4-6 per group) was topically treated with 3.2 mg NM dissolved in 200 μL acetone, while control mice received acetone alone. VD3 was dissolved in dimethyl sulfoxide, diluted in mineral oil, and administered via intraperitoneal injection (50 ng in 100 μL per mouse) 1 hour prior to NM exposure. Control mice received an equivalent volume of mineral oil. At 4 hours post-exposure, NM-treated skin was gently cleansed with 0.8% sodium hypochlorite followed by saline. Wound images were captured on days 1, 3, and 7. Mice were euthanized 7 days after NM exposure, and dorsal skin samples were collected and fixed in 10% neutral-buffered formalin for histological analysis as previously described.^40^

### 2.3 Statistical analysis

Quantitative data are presented as the mean ± standard deviation (SD). One-way or two-way analysis of variance (ANOVA) with Tukey’s or Bonferroni’s correction were used for multiple group comparisons. Unpaired two-tailed Student’s t-test was used for two different group comparisons. Pearson correlation was used to evaluate the associations. All statistical analyses were performed using R software (v4.0.0) and Prism 10.0 (GraphPad Software, USA). A PCvalue <C0.05 was considered statistically significant.

**Full descriptions of additional materials and methods are given in the Supporting Information.**

## 3. Results

### 3.1 NM induced dermal toxicity via remodeling the gut microbiota

In order to clarify the profile of the gut microbiota in NM-treated mice, we conducted metagenomic sequencing on fecal samples from mice in NM and control groups. The α-diversity (indexes of Shannon, Chao1, ACE, and Simpson) of NM-treated mice was not significantly different from that of the control group **(Fig. 1A and Fig. S1A-C)**. Notably, β-diversity analysis revealed a distinct clustering pattern in microbial communities between the two groups **(Fig. 1B)**. The analysis of similarities (ANOSIM) showed that the inter-group microbiota differences between NM-treated mice and controls were substantially greater than the intra-group differences **(Fig. 1C)**. At the species level, Venn diagrams identified 295 unique bacteria in controls, 194 in NM-treated mice, and 3815 shared by both **(Fig. S1D)**. Indicator analysis also revealed significant changes in the gut flora structure between the two groups **(Fig. 1D)**. Comparing the taxonomic composition of NM-treated mice to controls at the phylum, family, genus and species levels, we found that the relative abundance of *Proteobacteria, Verrucomicrobia*, *Escherichia, Parabacteroides, Akkermansiaceae*, *Akkermansia*, *Akkermansia muciniphila* (*A. muciniphila/AKK), Escherichia coli (E. coli)*, and *Muribaculaceae bacterium Isolate-105* was markedly decreased with the increase in *Mycoplasmataceae, Coriobacteriaceae, Pseudoflavonifractor, Enterococcus faecalis (E. faecalis),* and *Helicobacter apodemus* (**Fig. 1E and Fig. S1E-I**). Among which, *A. muciniphila* showed the largest reduction in abundance and was identified as the top contributor to group separation by the random forest model and variable importance in projection (VIP) score in partial least squares discrimination analysis (PLS-DA) **(Fig. 1F-H)**. Meanwhile, an receiver operating characteristic (ROC) analysis indicated that *A. muciniphila* was able to distinguish NM-treated mice from the controls with an area under the curve (AUC) value of 1, suggesting potential diagnostic value for NM-treated mice **(Fig. 1I)**. Accordingly, *A. muciniphila* was designated as the putative key functional microbe for further mechanistic studies of the gut microbiota in NM-induced dermal toxicity.

**Figure 1.**
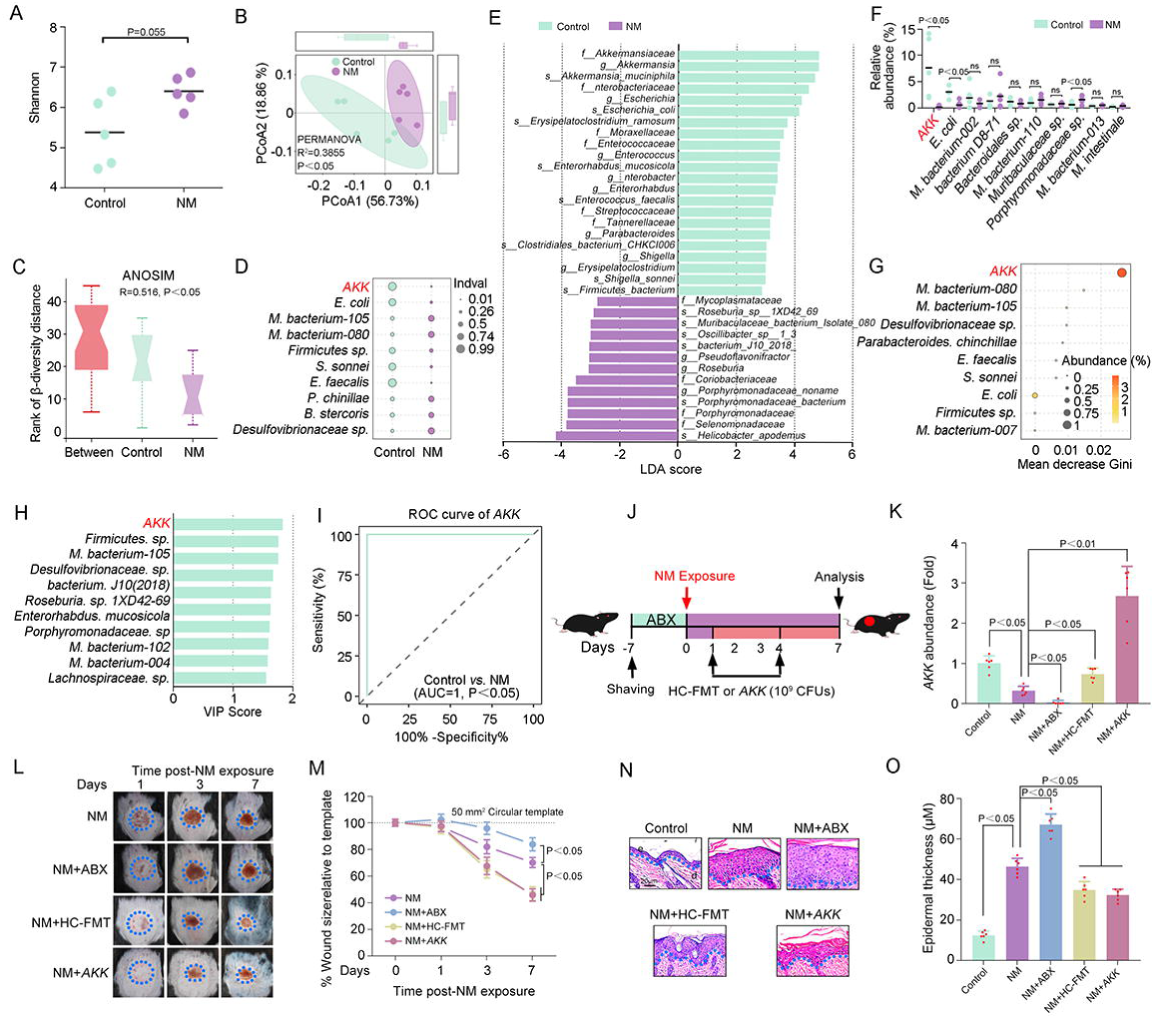
NM induced dermal toxicity via remodeling the gut microbiota. Six-week-old C57BL/6J mice (n = 5 per group) were exposed to NM on the dorsal skin. At 48 h post-exposure, fecal samples were collected for metagenomic analysis. (**A**) α-diversity assessed by the Shannon index of fecal microbiota in the control and NM groups. (**B**) PCoA based on Bray-Curtis distances, with the percentage of variation explained by each principal coordinate indicated in parentheses. (**C**) Comparison of β-diversity distances between the control and NM groups using ANOSIM. (**D**) Indicator value analysis (Indval) identifying species significantly enriched in each group. (**E**) Linear discriminant analysis (LDA) scores derived from LEfSe analysis, showing the biomarker taxa (LDA score >2 and a significance of P<0.05 determined by the Wilcoxon signed-rank test. (**F)**The relative abundance of the top 10 enriched taxa in the control and NM groups. (**G**) Variable importance ranked by the Gini index using the random forest algorithm. (**H**) VIP scores derived from OPLS-DA. (**I**) ROC curve demonstrating the discriminative ability of *AKK*. (**J**) Schematic overview of the experimental design. (**K**) *AKK* abundance measured by qPCR (n = 6 per group). (**L**) Representative images of skin injury at 1, 3, and 7 days post NM exposure. (**M**) Quantification of wound size corresponding to (L) (n = 6 per group). Mice were sacrificed at 7 days post NM exposure, and dorsal skin tissues were collected. (**N**) H&E staining for assessment of epidermal thickness. (**O**) Quantification of the epidermal thickness related to (N) (n = 6 per group). Data are expressed as mean ± SD. **See also Figure S1**.

To further identify the exact role of the gut microbiota in NM-caused dermal toxicity, NM-exposed mice were subjected to broad-spectrum antibiotics (ABX) treatment, fecal samples from the controls transplantation (HC-FMT), or *A. muciniphila* monocolonization, respectively **(Fig. 1J)**. In line with previous observations,^45^ ABX resulted in significant reduction in *A. muciniphila* abundance, meanwhile, HC-FMT and *A. muciniphila* markedly increased the abundance of *A. muciniphila* in NM-treated mice (**Fig. 1K)**. As shown in **Fig. 1L-O**, ABX significantly increased the size of damaged-skin area and epidermal thickness in NM-exposed mice; in contrast, HC-FMT or *A. muciniphila* supplementation markedly attenuated NM-induced skin injury. These collective data indicated that the gut microbiota played a pivotal role in regulating NM-caused skin injury, among which *A. muciniphila* (a well established SCFA-producing probiotic) emerged as the central functional microbe.

### 3.2 Gut microbiota-derived BA attenuated NM-caused dermal toxicity

A growing body of evidence indicates that the metabolites of the gut microbiota, especially SCFAs, are key mediators of microbiota-gut-skin communication ^17^. Thus, targeted metabolomics was employed to detect the metabolites in feces and serum from NM-treated mice or the controls. The metabolic profiles of the feces were different between the two groups **(Fig. 2A)**. A total of 205 metabolites were characterized, including 40 amino acids (31.30% of total), 10 SCFAs (26.87%) and 15 carbohydrates (12.26%) **(Fig. 2B)**. Compared to those in the controls, the contents of 10 metabolites (VIP ≥ 1, |log_2_FC | ≥ 1.5, and *p* value < 0.05, classified as amino acids, SCFAs, bile acids and so on) were notably changed in feces of NM-treated mice **(Fig. 2C)**. Depending on the random forest model, support vector machine (SVM), Boruta and Venn diagram analyses, BA and isobutyric acid were found to be the potential metabolites for separating NM-treated mice from the controls, among which fecal BA was the top metabolite in the group separation **(Fig. 2D-G)**. Concurrently, we detected significant reductions in fecal acetic acid, propionic acid, and BA levels in NM-treated mice, with BA exhibiting the most pronounced decrease **(Fig. 2H)**. The ROC analysis revealed that fecal BA could be used to distinguish NM-treated mice from the controls with an AUC value of 1, suggesting potential diagnostic value for NM-treated mice **(Fig. 2I)**. We also found that fecal BA contents were positively correlated with the abundance of *Akkermansia* and *A. muciniphila* **(Fig. 2J and K)**. Moreover, we identified that the levels of serum propionic acid were not significantly changed; while, serum acetic acid and BA contents were notably reduced in NM-treated mice **(Fig. 2L)**. The ROC analysis revealed that serum BA could also distinguish NM-treated mice from the controls (AUC = 1) **(Fig. 2M)**. Serum BA content was also positively correlated with the abundance of *A. muciniphila* **(Fig. 2N)**. These results indicated that BA reduction represents a signature characteristic of the microbial metabolome in NM-treated mice, suggesting its critical role in NM-induced dermal toxicity.

**Figure 2.**
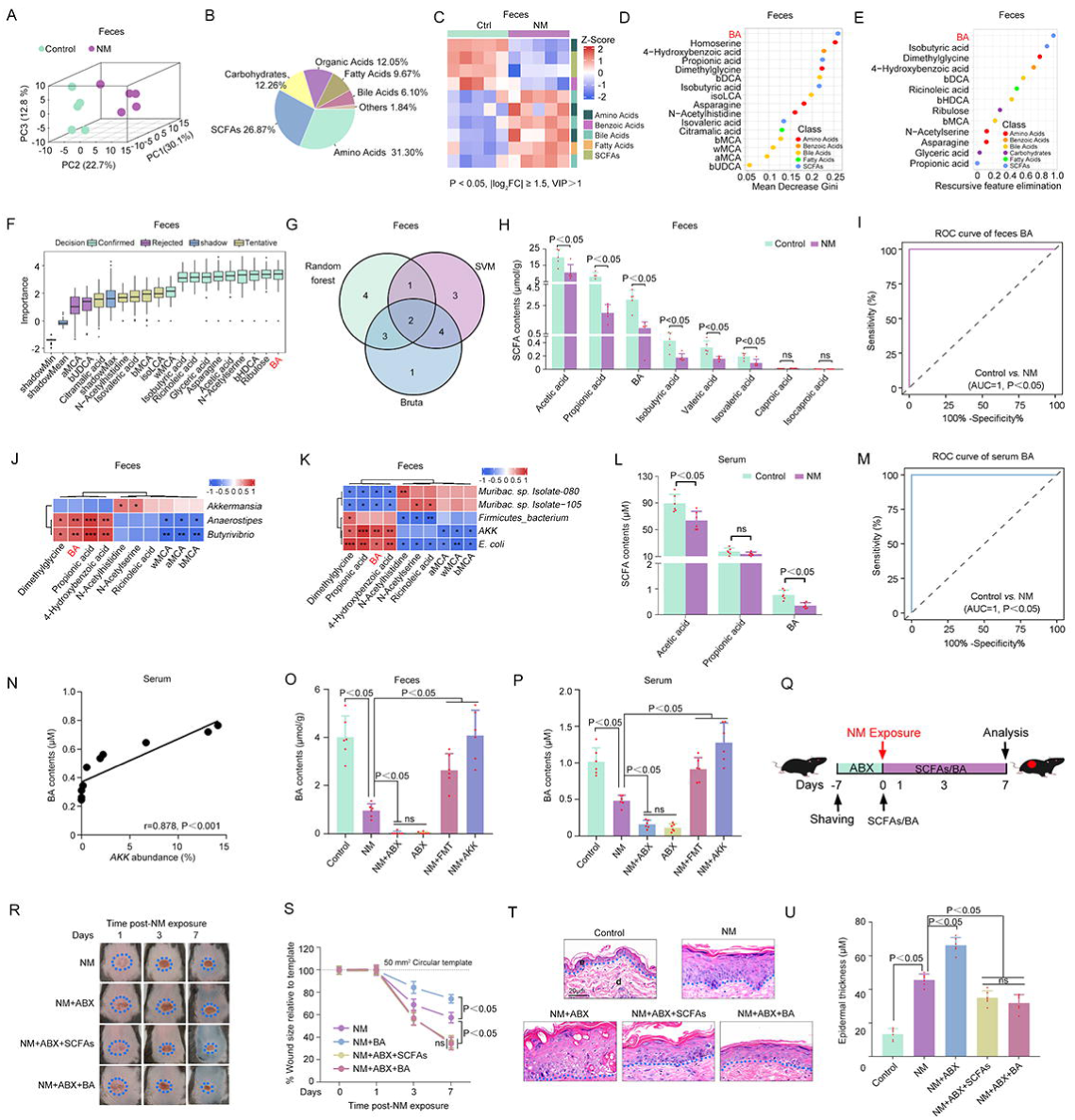
Gut microbiota-derived BA attenuated NM-caused dermal toxicity. Targeted metabolomic analysis was performed on fecal samples collected from mice (n = 5 per group) at 48 h following dorsal skin exposure to NM. (**A**) OPLS-DA of metabolite profiles in fecal samples from control and NM-treated mice. (**B**) Pie chart illustrating the compositional distribution of identified metabolites. (**C**) Heatmap showing the differential abundant metabolites in NM group compared with the control group. Metabolites were selected based on the following criteria: P < 0.05, |log_2_FC| > 1.5, and VIP > 1. Ranking of metabolite importance using **(D)** Random forest, **(E)** SVM, and **(F)** Boruta. (**G**) Venn diagram showing the number of common metabolites identified by all three machine learning algorithms. (**H**) Fecal SCFA concentrations in the control and NM groups. (**I**) ROC curve analysis of BA in fecal samples. Heatmap of the Pearson correlation between fecal metabolite contents and the gut microbiota abundance at the **(J)** family and (K) species levels. Red indicates a positive association, blue a negative association. The statistical significance is denoted on the squares (*P < 0.05, **P < 0.01 and ***P < 0.001). **(L)** Serum SCFA concentrations in the control and NM groups (n = 6 per group). (**M**) ROC curve analysis of BA in serum. (**N**) Pearson correlation between serum BA levels and *AKK* abundance (n = 6 per group). (**O**) Comparison of fecal BA levels among indicated groups (n = 6 per group). (**P**) Comparison of serum BA levels among indicated groups (n = 6 per group). (**Q**) Schematic diagram of the experimental workflow. (**R**) Representative images of skin wound at days 1, 3, and 7 after NM exposure. (**S**) Quantification of the wound size related to (R) (n = 6 per group). Mice were sacrificed at 7 days post NM exposure, and dorsal skin tissues were collected. (**T**) H&E staining for assessment of epidermal thickness. (**U**) Quantification of the epidermal thickness related to (T) (n = 6 per group). Data are expressed as mean ± SD; ns: no significance.

As shown in **Fig. 2O and P**, ABX notably decreased BA contents, but HC-FMT or *A. muciniphila* markedly increased BA levels in the serum and feces of NM-treated mice. Meanwhile, a mixture of SCFAs (67.5 mM acetate, 25.9 mM propionate, and 40 mM BA) or 100 mM BA in the drinking water was used to directly investigate the effect of SCFAs or BA on NM-caused skin injury **(Fig. 2Q)**. As expected, BA or SCFA mix significantly attenuated ABX-caused increase of skin wound size and epidermal thickness in NM-treated mice; these indices were not different between BA- and SCFA mix-treated groups **(Fig. 2R-U)**. Collectively, these findings suggested that the gut microbiota (especially *A. muciniphila*)-generated BA could serve as the potential functional inhibitor of NM-induced skin injury.

### 3.3 NM decreased microbial BA levels by inhibiting α-defensin production

To identify the potential mechanisms that contribute to the effect of NM on microbial BA production, RNA-Seq analysis was performed on SI tissue from NM-treated and control mice. As shown in **Fig. 3A**, NM significantly altered the transcriptome of SI tissue in mice, resulting in 180 downregulated genes and 88 upregulated genes (|log_2_FC| ≥ 1.0 and *p* < 0.05). Gene Ontology (GO) enrichment analyses identified pathways that were associated with antimicrobial humoral response, antimicrobial humoral response mediated by antimicrobial peptide and defense response to bacterium **(Fig. 3B)**. Among these, the pathway of defense response to bacterium had the most differentially expressed genes (DEGs) (38/171) **(Fig. 3B)**. The differential expression patterns of these 38 DEGs in this functional category contained 20 α-defensin genes (*Defa3, Defa5, Defa20, Defa21, Defa22, Defa23, Defa24,* etc) and 6 genes related to the regulation of α-defensin function (*Ang4, Lyz1, Reg3*γ*, Nod2, Gsdmc2*, and *Gsdmc3*) **(Fig. 3C)**, indicating that α-defensins might play an important role in NM’s regulation of the gut microbiota and BA generation.

**Figure 3.**
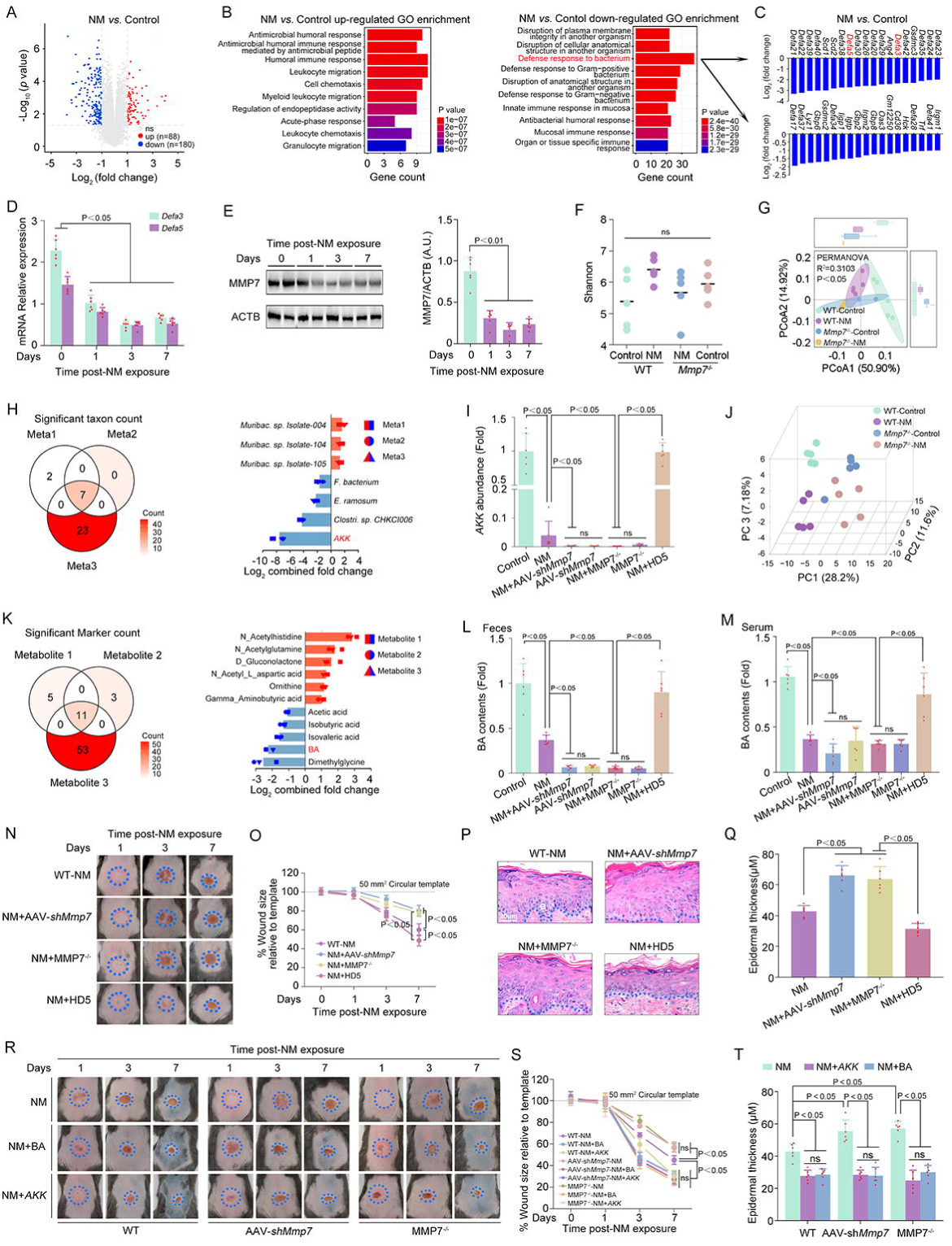
NM decreased microbial BA levels by inhibiting α-defensin production. At 48 h after NM exposure, mice from each group (n = 4 per group) were sacrificed, and a 0.5 cm segment of the terminal ileum (distal to the cecum) was excised and subjected to transcriptomic profiling. (**A**) Volcano plot showing DEGs between the control and NM groups. (**B**) GO enrichment analysis of upregulated (left) and downregulated (right) DEGs, respectively. (**C**) Bar plot displaying the most significantly downregulated genes in NM group compared with the control group. (**D**) Relative expression of *Defa3* and *Defa5* mRNA at different time points after NM exposure (n = 6 per group). (**E**) Western blot analysis of MMP7 levels at indicated time points post-NM exposure and the bar graphs show the quantification of MMP7 (n = 6 per group). **(F)** Comparison of the Shannon index between WT and MMP7*^−/−^*mice (n = 5 per group). (**G**) PCoA based on Bray-Curtis distances of fecal microbial composition in WT and MMP7*^−/−^* mice (n = 5 per group). (**H**) Meta-analysis of metagenomic data identifying consistently altered microbial taxa across datasets (n = 5 per group). (**I**) *AKK* abundance measured by qPCR. (**J**) OPLS-DA of metabolite profiles in WT and MMP7*^−/−^* mice (n = 5 per group). (**K**) Meta-analysis of metabolites revealing concordant changes in metabolite levels (n = 5 per group). BA levels in the indicated groups in **(L)** fecal and **(M)** serum samples (n = 6 per group). **(N)** Representative images of skin injury at 1, 3, and 7 days after NM exposure. **(O)** Quantification of the wound size related to (N) (n = 6 per group). **(P)** H&E staining for assessment of epidermal thickness. **(Q)** Quantification of the epidermal thickness related to (P) (n = 6 per group). **(R)** Representative images of skin wounds at 1, 3, and 7 days after NM exposure. **(S)** Quantification of the wound size related to (S) (n = 6 per group). (**T**) Quantification of the epidermal thickness for H&E staining assessment (n = 6 per group). Data are presented as mean ± SD; A.U.: arbitrary units; ns: no significance. **See also Figure S2**.

Quantitative polymerase chain reaction (qPCR) analysis further revealed that the expression of *Defa3* and *Defa5* (the most important and effective α-defensins in C57BL/6J mice) was significantly decreased in NM-treated mice **(Fig. 3D)**. It has been demonstrated that MMP7 is involved in α-defensin maturation in mice.^46^ As expected, we found that NM inhibited MMP7 expression in SI tissue **(Fig. 3E)**. Meanwhile, adenovirus-associated virus serotype 9 carrying with a *Mmp7* short hairpin RNA (AAV-*shMmp7*), MMP7^−/−^ mice and MMP7^−/−^;Tg(HD5) mice were used to further define the role of α-defense in NM’s effect on the gut microbiota and BA production. As shown in **Fig. S2A and B**, AAV-*shMmp7* and MMP7^−/−^ mice significantly inhibited MMP7 expression and HD5 overexpression notably increased *Hd5* expression in SI tissue. Moreover, the exact role of α-defensins in NM-induced alteration of the gut microbiota was directly investigated by metagenomic sequencing analyses. As shown in **Fig. 3F and G**, bacterial species alpha diversity showed no significant differences among all samples, beta diversity analysis revealed distinct microbiota features in the control group relative to NM-treated wild type (WT) mice or MMP7^−/−^ mice; while, the structure of the gut microbiota was similar in MMP7^−/−^mice regardless of NM treatment. To gain further insight into the role of α-defensins in NM-induced bacterial species modulation, we combined three meta-analyses of different comparisons across experiments: Meta1, NM (WT-control *vs.* WT-NM)-induced differences in line with those driven by genotype (WT-control *vs.* MMP7^−/−^ - control); Meta 2, NM (WT-control *vs.* WT-NM) -induced differences in line with those driven by NM+genotype (WT-control *vs.* MMP7^−/−^-NM); Mate 3, genotype (WT-control *vs.* MMP7^−/−^-control)-induced differences in line with those driven by NM+genotype (WT-control *vs.* MMP7^−/−^-NM) **(Fig. S2C-E)**. This approach identified 7 consistently changed taxa, with *A. muciniphila* abundance showing the greatest reduction **(Fig. 3H)**. Both NM exposure and MMP7 knockdown notably reduced *A. muciniphila* levels, with no additional decrease seen in NM-treated MMP7-deficient mice **(Fig. 3I)**. In contrast, HD5 overexpression increased *A. muciniphila* abundance in NM-treated MMP7^−/−^;Tg(HD5) mice **(Fig. 3I)**. These data suggested that α-defensins played a pivotal role in NM-induced gut microbiota modulation, especially in reducing *A. muciniphila* abundance.

Finally, the role of α-defensins in NM-induced BA reduction was also determined. As shown in **Fig. 3J**, the metabolic profiles of the feces were differed between the control group and both NM-treated mice and MMP7^−/−^ mice; however, MMP7^−/−^ mice showed similar metabolite composition in the presence or absence of NM. Integration of three meta-analyses across experimental comparisons as detailed above (Mate 1-3) was also performed to further clarify the involvement of α-defensins in NM-induced BA decline **(Fig. S2F-G)**. This approach identified 11 consistently changed metabolites across conditions, among which BA levels exhibited the second most pronounced decline **(Fig. 3K)**. BA levels were substantially reduced by either NM exposure or MMP7 knockdown, but no further reduction was observed in NM-treated MMP7-deficient mice **(Fig. 3L and M)**. In contrast, HD5 overexpression restored BA levels in NM-treated MMP7^−/−^;Tg(HD5) mice **(Fig. 3L and M)**. In addition, we found that MMP7 deficiency promoted NM-caused skin injury, which was attenuated by HD5 overexpression, *A. muciniphila* transplantation, or BA supplementation **(Fig. 3N-T and Fig. S2I)**. These data collectively confirmed that NM remodeled the gut microbiota particularly reducing *A. muciniphila* abundance by inhibiting α-defensin generation, thereby reducing BA contents, ultimately accelerating skin injury.

### 3.4 NM suppressed α-defensin production via inhibiting intestinal VDR

Next, the potential involvement of VDR in NM-induced α-defensin decline was investigated. We found that NM significantly inhibited VDR expression in SI tissue **(Fig. 4A)**. Meanwhile, adenovirus-associated virus serotype 9 carrying with a *Vdr* short hairpin RNA (AAV-*shVdr*) was applied to further identify the role of VDR in NM’s effect on α-defensin production. As shown in **Fig. 4B and C**, AAV-*shVdr* notably knocked down VDR expression in SI tissue; however, NM did not additionally suppress MMP7, *Defa3,* and *Defa5* expression in VDR-silenced mice. Moreover, NM also failed to further decrease *A. muciniphila* abundance and BA levels in the presence of AAV-*shVdr* in mice **(Fig. 4D-F)**. Finally, VDR knockdown in SI tissue markedly exacerbated NM-caused skin injury **(Fig. 4G-J)**. Our data suggested that intestinal VDR was required for NM-induced inhibition of α-defensin generation, which subsequently reduced *A. muciniphila* abundance and BA levels, ultimately promoting dermal toxicity.

**Figure 4.**
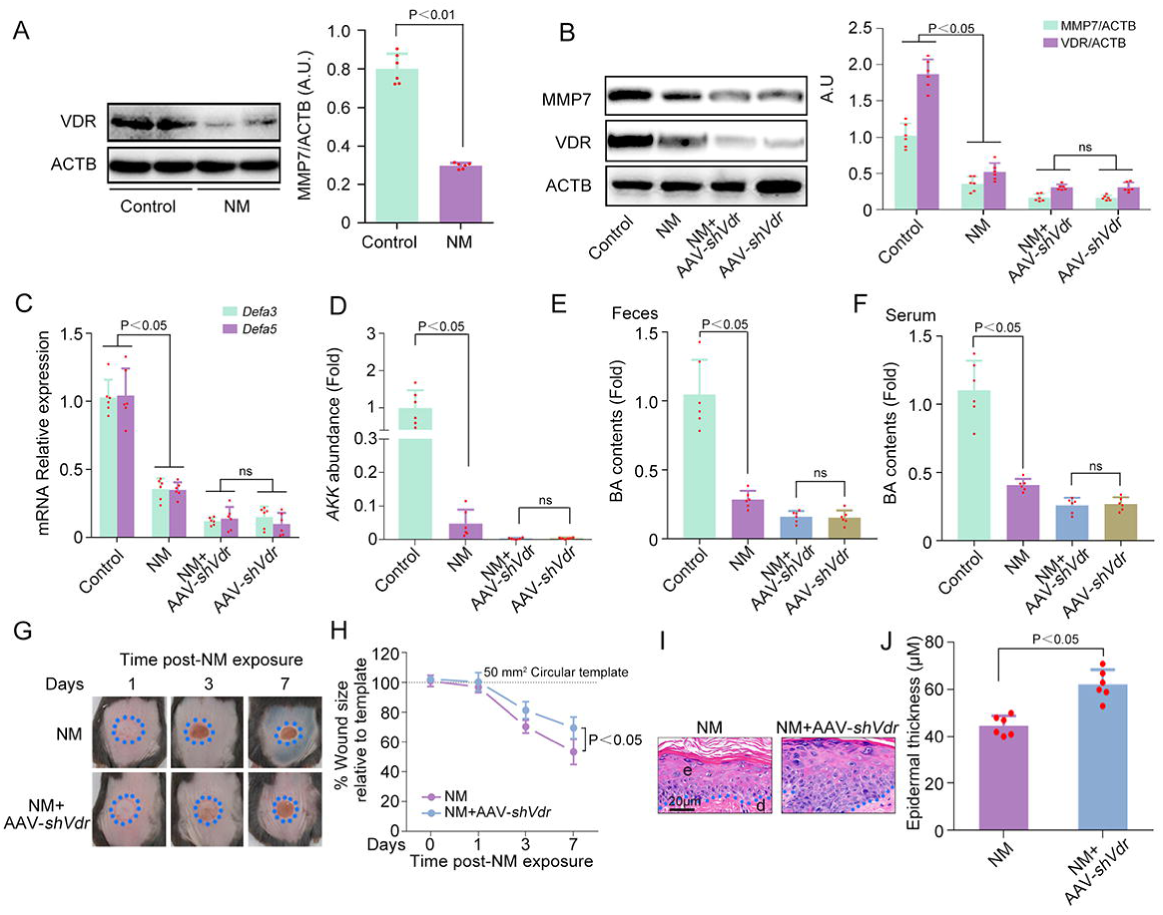
NM suppressed α-defensin production via inhibiting intestinal VDR. At 48 h post-NM exposure on the dorsal skin of mice, ileal tissues were collected for analysis. (**A**) Western blot analysis of VDR expression with quantification shown in the bar graphs. **(B)** The expression of VDR and MMP7 was detected by western blot and the bar graphs show the quantification of the indicated proteins. (**C**) Relative expression of *Defa3* and *Defa5* mRNA determined by qPCR. (**D**) *AKK* abundance detected by qPCR. BA concentrations in **(E)** fecal and **(F)** serum samples from mice with indicated treatments. **(G)** Representative images of skin wounds at indicated time points after NM exposure. **(H)** Quantification of the wound size related to (G). **(I)** H&E staining for assessment of epidermal thickness. **(J)** Quantification of the epidermal thickness related to (I). Data are presented as mean ± SD (n = 6 per group); A.U.: arbitrary units; ns: no significance.

### 3.5 VD3 attenuated NM-caused dermal toxicity by reshaping the gut microbiota

Reportedly, VD3 has significant protective effect against NM-caused dermal toxicity;^40^ however the exact underlying mechanisms need to be elucidated. Based on our above findings, we investigated the role of the gut microbiota in VD3’s beneficial effect against NM-induced skin injury. In agreement with previous published data,^40^ VD3 promoted wound healing and enhanced new epidermis maturation in NM-exposed skin **(Fig. 5A-D)**. Metagenomic sequencing analysis revealed that the α diversity (indexes of Shannon and Simpson) was decreased in VD3-treated NM-exposed mice, indicating the loss of particular bacterial taxa **(Fig. 5E and Fig. S3A)**. Principal coordinates analysis (PCoA) of Bray-Curtis distance found obvious differences in the gut microbial composition between the two groups **(Fig. 5F)**. ANOSIM demonstrated greater microbiota dissimilarity between NM and NM+VD3 treatment groups than within each group **(Fig. 5G)**. At the species level, Venn diagrams identified 689 unique bacteria in NM group, 722 in NM+VD3-treated mice, and 4164 shared by both **(Fig. S3B)**. Indicator analysis also revealed significant changes in the gut flora structure between the two groups **(Fig. 5H)**. Compared to those in NM group, the relative abundance of *Verrucomicrobia, Akkermansiaceae*, *A. muciniphila,* and *E. coli* was markedly increased by VD3 in NM-treated mice **(Fig. 5I and Fig. S3C-F)**. Of these, *A. muciniphila* exhibited the most pronounced decrease in abundance and was identified as the primary species contributing to group separation by the random forest model and VIP score in PLS-DA **(Fig. 5J-L)**. Meanwhile, ROC analysis demonstrated *A. muciniphila* as an excellent biomarker (AUC = 1) for distinguishing NM+VD3-treated mice from NM-treated ones **(Fig. 5M)**. Furthermore, the gut microbiota depletion by ABX abolished the beneficial effect of VD3 on NM-caused dermal toxicity, which was reversed by *A. muciniphila* **(Fig. 5N-Q)**. These results indicated that VD3 remodeled the gut microbiome especially enriching *A. muciniphila*, thereby ameliorating NM-induced skin injury.

**Figure 5.**
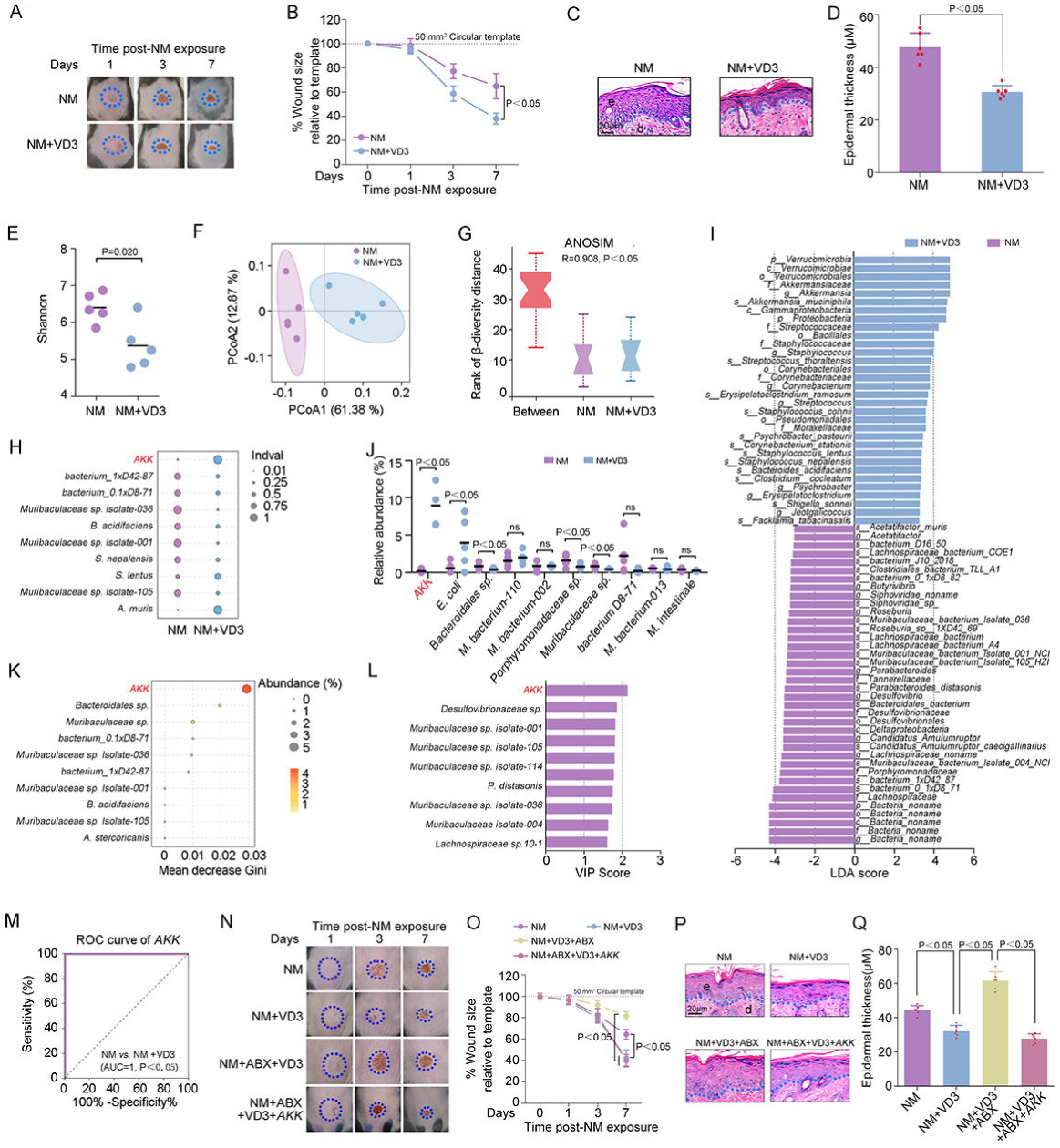
VD3 attenuated NM-caused dermal toxicity by reshaping the gut microbiota. Mice (n = 6 per group) received dorsal skin exposure to NM with or without VD3 treatment (50 ng/100 μL per mouse, i.p.) as described in the Materials and Methods. **(A)** Representative images of dorsal skin lesions in NM and NM + VD3 groups at days 1, 3, and 7 post-exposure. **(B)** Quantitative analysis of the wound size related to (A). (**C**) Representative H&E-stained sections of dorsal skin. **(D)** Quantification of the epidermal thickness related to (C). Fecal samples from the indicated groups were collected and processed for metagenomic sequencing (n = 5 per group). (**E**) α-diversity measured by the Shannon index in NM and NM + VD3 groups. **(F)** PCoA of β diversity using Bray-Curtis metric distance with PERMANOVA significance test. **(G)** Similarities of (F) were analyzed by ANOSIM test. (**H**) Indval revealing significantly enriched taxa in each group. (**I**) LDA scores derived from LEfSe analysis, showing the biomarker taxa (LDA score >2 and a significance of P <0.05 determined by the Wilcoxon signed-rank test). (**J**) The relative abundance of the top 10 differentially abundant taxa between NM and NM + VD3 groups. (**K**) Variable importance ranking based on the Gini index calculated by random forest algorithm. (**L**) VIP scores of taxa derived from OPLS-DA. (**M**) ROC curve evaluating the discriminative ability of *AKK*. (**N**) Representative images of skin wounds in the indicated groups at 1, 3, and 7 days after NM-exposure. **(O)** Quantification of the wound size related to (N) (n = 6). (**P**) H&E staining was performed for analysis of epidermal thickness. **(Q)** Quantification of the epidermal thickness related to (P) (n = 6). Data are presented as mean ± SD; ns: no significance. **See also Figure S3**.

### 3.6 Microbial BA mediated the beneficial effect of VD3 on NM-caused dermal toxicity

According to our above results, the role of gut microbiota-produced BA in the beneficial effect of VD3 was further explored. Targeted metabolomics analysis found that the metabolic profile of feces was different in NM and NM+VD3 groups **(Fig. 6A and B)**. Among which, BA was recognized as the top metabolite in feces that could be applied to discriminate VD3+NM from NM group by random forest model analysis **(Fig. 6C)**. ROC analysis confirmed BA as a robust biomarker for distinguishing NM+VD3-treated mice from NM-treated ones, with an AUC of 1 **(Fig. 6D)**. Meanwhile, BA contents were notably increased in feces and serum of NM+VD3-treated mice, which were positively correlated with *A. muciniphila* abundance **(Fig. 6E-H)**. We also found that *A. muciniphila* markedly reversed ABX-mediated decline in BA levels in both feces and serum of NM+VD3-treated mice **(Fig. 6I and J)**. Finally, we identified that BA significantly attenuated ABX-caused inhibition of the protective effect of VD3 against NM-caused skin injury **(Fig. 6K-N)**. These findings suggested that BA could be the potential functional metabolite produced by the gut microbiota, especially *A. muciniphila,* which could mediate the protective effect of VD3 against NM-caused dermal toxicity.

**Figure 6.**
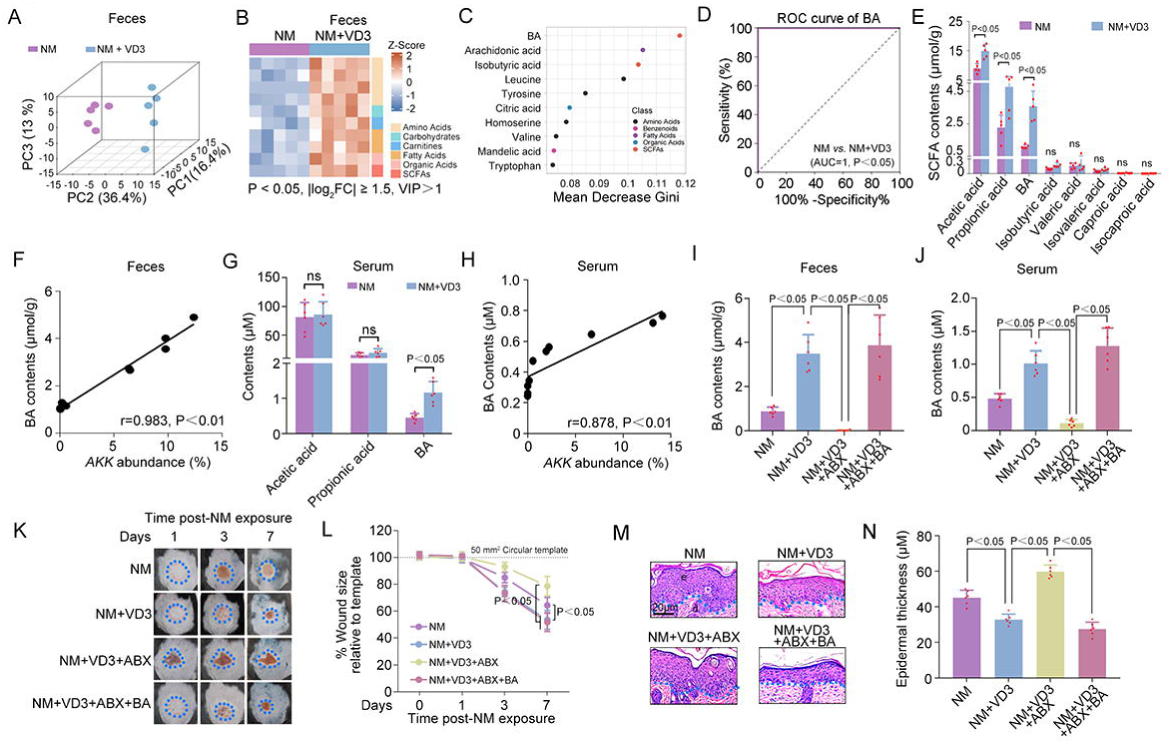
Microbial BA mediated the beneficial effect of VD3 on NM-caused dermal toxicity. Fecal metabolomic analysis of mice in the indicated groups after dorsal skin NM exposure with or without VD3 treatment. (**A**) OPLS-DA of fecal metabolite profiles in NM and NM + VD3 groups (n = 5 per group). (**B**) Heatmap displaying the top 10 differentially abundant metabolites based on the following criteria: *P* < 0.05, |log_2_FC| > 1.5, VIP > 1 (n = 5 per group). (**C**) Ranking the importance of differential metabolites through Gini index. (**D**) ROC curve evaluating the discriminatory power of fecal BA between NM and NM + VD3 groups (n = 5 per group). (**E**) Fecal SCFA levels in NM and NM+VD3 groups (n = 5 per group). (**F**) Pearson correlation between fecal BA levels and *AKK* abundance (n = 5 per group). (**G**) Serum SCFA levels in NM and NM+VD3 groups (n = 6 per group). (**H**) Pearson correlation between serum BA levels and *AKK* abundance (n = 5 per group). BA levels in **(I)** fecal and **(J)** serum samples across indicated groups (n = 6 per group). **(K)** Representative images of skin wounds in NM and NM + VD3 groups at days 1, 3, and 7 post-exposure. **(L)** Quantification of the wound size related to (K) (n = 6 per group). **(M)** H&E staining was performed for analysis of epidermal thickness. **(N)** Quantification of the epidermal thickness related to (M) (n = 6). Data are presented as mean ± SD; ns: no significance. .

### 3.7 VD3 increased microbial BA levels via the intestinal VDR-α-defensin signaling pathway

Our current study identified that VDR-mediated α-defensin generation plays a core role in NM-induced microbial BA reduction and subsequent dermal toxicity. Accordingly, the potential relationship between VD3 and VDR-mediated α-defensin generation was investigated in NM-treated mice. As shown in **Fig. 7A-B and Fig. S4A,** VD3 increased *Defa3*, *Defa5*, and MMP7 expression in NM-exposed WT mice. Meanwhile, bacterial species alpha diversity showed no significant differences among all samples **(Fig. S4B)**. Beta diversity analysis revealed distinct microbiota features in NM-treated WT mice relative to the control- or VD3+NM-treated WT mice; the microbiota composition of VD3+NM-treated MMP7^−/−^ mice was different from that of VD3+NM-treated WT mice, which resembled that of the control WT mice; however, the gut microbiota composition was similar in MMP7^−/−^mice regardless of treatments **(Fig. S4C)**. To gain further insight into the role of α-defensins in VD3-induced bacterial species modulation in NM-treated mice, we combined three meta-analyses of different comparisons across experiments: Meta1, differences driven by NM (WT-control *vs.* WT-NM) consistent with those driven by VD3 (WT-NM *vs.* WT-NM+VD3); Meta 2, differences driven by VD3 (WT-NM *vs.* WT-NM+VD3) consistent with those driven by VD3+genotype (MMP7^−/−^-NM *vs.* WT-NM+VD3); Mate 3, differences driven by genotype (MMP7^−/−^-VD3+NM *vs.* WT-NM+VD3) consistent with those driven by VD3+genotype (MMP7^−/−^-NM *vs.* WT-NM+VD3) **(Fig. S4D-F)**. This approach identified 4 consistently changed taxa, among these only *A. muciniphila* abundance was increased **(Fig. 7C)**. VD3 significantly increased *A. muciniphila* levels in NM-treated mice, an effect that was abolished by MMP7 deficiency **(Fig. 7D)**. These results indicated that α-defensin was involved in VD3-induced gut microbiota modulation particularly in enriching *A. muciniphila*.

**Figure 7.**
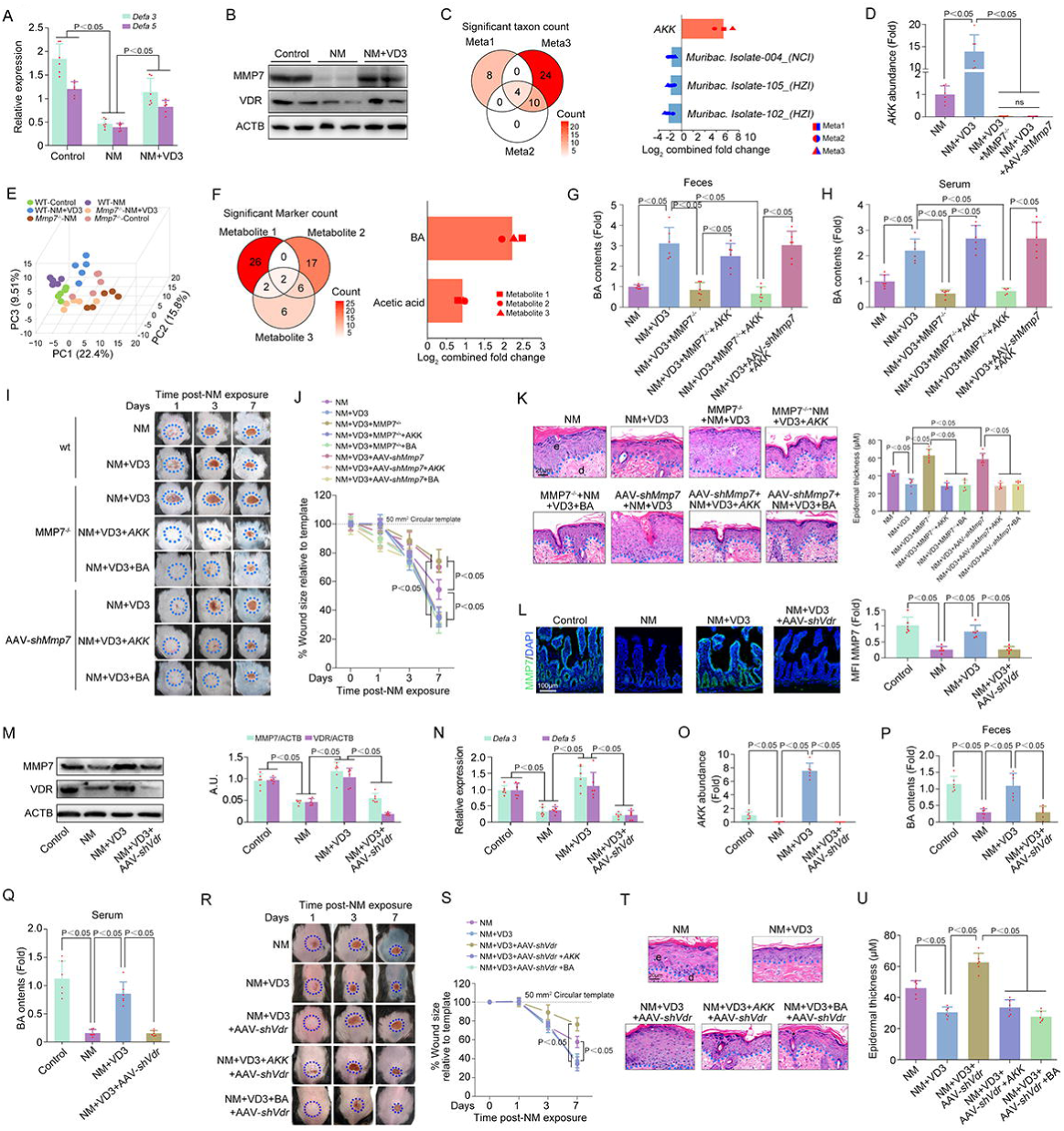
VD3 increased microbial BA levels via the intestinal VDR-α-defensin signaling pathway. (**A**) Relative expression of *Defa3* and *Defa5* mRNA (n = 6 per group). (B) Western blot analysis of VDR and MMP7 expression. (**C**) Meta-analysis of metagenomic sequencing data identifies microbial taxa consistently altered in the same direction across datasets (n = 5 per group). (**D**) Relative abundance of *AKK* across the indicated groups. (**E**) OPLS-DA of fecal metabolite profiles in the indicated groups (n = 5 per group). (**F**) Meta-analysis of targeted metabolomics data identifying metabolites consistently altered in the same direction across datasets (n = 5 per group). BA levels in **(G)** fecal and **(H)** serum samples from the indicated groups (n = 6 per group). **(I)** Representative images of dermal lesions at 1, 3, and 7 days post NM exposure. **(J)** Quantification of the wound size related to (I) (n = 6 per group). (**K**) Representative H&E-stained sections and corresponding quantification of epidermal thickness. (**L**) Immunofluorescence staining and the quantification of MMP7 in ileal villus epithelium across the indicated groups (n = 6 per group). (**M**) Western blot was used to detect the levels of MMP7, VDR, and ACTB proteins expression and the bar graphs show the quantification of the indicated proteins (n = 6 per group). (**N**) Relative expression of *Defa3* and *Defa5* mRNA across the indicated groups (n = 6 per group). (**O**) *AKK* abundance detected by qPCR (n = 6 per group). BA levels in **(P)** fecal and **(Q)** serum samples across the indicated groups (n = 6 per group). (**R**) Representative images of skin wounds at 1, 3, and 7 days post NM exposure. **(S)** Quantification of the wound size related to (R) (n = 6 per group). **(T)** H&E staining was performed for analysis of epidermal thickness. **(U)** Quantification of the epidermal thickness related to (T) (n = 6 per group). Data are presented as mean ± SD; A.U.: arbitrary units. **See also Figure S4**.

Furthermore, the metabolic profiles of the feces were differed in NM-treated WT mice relative to the control- or VD3+NM-treated WT mice; the metabolite composition of VD3+NM-treated MMP7^−/−^ mice was different from that of VD3+NM-treated WT mice, which resembled that of the control WT mice; however, the composition was similar in MMP7^−/−^ mice regardless of treatments **(Fig. 7E)**. Integration of three meta-analyses across experimental comparisons as detailed above (Meta 1-3) was also performed to further clarify the involvement of α-defensins in VD3-induced BA generation **(Fig. S4G-I)**. 2 consistently altered metabolites across conditions were identified, among which BA levels exhibited a more substantial increase **(Fig. 7F)**. The VD3-induced increase in BA levels in NM-treated mice was inhibited by MMP7 knockdown, an effect that was reversed by *A. muciniphila* supplementation **(Fig. 7G and H)**. Additionally, MMP7 knockdown abolished VD3’s protective effect against NM-caused skin injury, which was attenuated by *A. muciniphila* transplantation or BA supplementation **(Fig. 7I-K)**. Finally, we found that the protective effects of VD3 against NM-caused dermal toxicity, decreases in α-defensin contents, MMP7 expression, *A. muciniphila* abundance, and BA levels were abolished by AAV-*shVdr* in mice, effects that were rescued by *A. muciniphila* or BA supplementation (**Fig. 7L-U**). These data collectively implicated that the intestinal VDR-α-defensin signaling contributed to VD3-mediated microbial BA production and the subsequent mitigation of NM-induced dermal toxicity.

## 4. Discussion

In the present study, we identified for the first time that NM caused dermal toxicity through remodeling the gut microbiota. The fact that the gut microbiota is related to skin disorders has been increasingly recognized. Previous studies have shown that compared to healthy mice, differences in the α and β diversity, increase abundance of *Firmicutes* and decrease of *Bacteroidota* are observed in a mouse model of UVB-induced skin damage.^47^ The microbial ecosystem in diabetic patients has less diversity than that in healthy controls, whereas certain opportunistic pathogens such as *Clostridium hathewayi, clostridium symbioum,* and *E. coli*, are overrepresented in the diabetic group, accompanied by depletion of SCFA-producing bacteria, a condition closely associated with chronic diabetic wounds.^48,49^ In 2024, Dokoshi et al. confirmed that disruption of the dermis by giving a 1.5 cm full-thickness incisional wound results in gut microbiota dysbiosis thereby influencing susceptibility to disease in mice, providing direct evidence of a skin-gut axis by demonstrating that damage to the skin alters the gut microbiome.^50^ White and his colleagues first showed that depletion of the gut microbiota attenuates the delayed lethal effect of NM in mice, supporting the hypothesis that the gut flora is involved in the pathogenesis of NM-mediated systemic toxicity.^51^ However, the exact characteristic of NM-altered gut microbiota, especially the core microbes, is not well defined. Herein, we uncovered that a decrease in the abundance of SCFA-producing bacteria, especially *A. muciniphila,* was a hallmark feature of the gut microbiota in NM-exposed mice. Moreover, depletion of the gut microbiota by ABX significantly promoted NM-caused skin injury; while, HC-FMT or *A. muciniphila* notably ameliorated NM-induced cutaneous damage in mice. These findings suggested that NM induced dermal toxicity via modulating the gut microbiota and that *A. muciniphila* represented a potential therapeutic probiotic. Recently, oral administration of *Bifidobacterium longum* or *Lactobacillus plantarum HY7714* has been found to notably alleviate UVB-induced photoaging.^25,52^ And a multi-strain probiotic formulation comprising *Lactobacillus acidophilu*s, *Lactobacillus casei*, *Lactobacillus fermentum*, and *Bifidobacterium bifidum* substantially alleviates ulcer metrics (length, width, and depth) in patients with diabetic foot ulcers.^53^ These data indicate that the probiotic supplementation is a potentially effective therapy for skin wound healing. Our findings complement previous studies on the role of the gut microbiota in skin disorders especially wound healing, indicating that the intestinal flora particularly *A. muciniphila,* is a valuable tool for NM-caused skin injury.

Microbiota-generated SCFAs, mainly containing acetate, propionate, and BA, are considered vital metabolites that contribute to the relationship between the intestinal flora and diseases, including skin disorders and injuries.^54,55^ Clinical studies have demonstrated that BA levels are lower in chronic spontaneous urticaria patients than those in healthy controls, which are negatively correlated with the development of dermatitis.^56,57^ *Bifidobacteria adolescentis* ameliorates DNFB-induced atopic dermatitis by enhancing BA generation, which concomitantly drives Treg differentiation and inhibits Th2 responses in mice.^58^ Meanwhile, supplementation with acetate and propionate significantly attenuates imiquimod-induced psoriasis in mice.^21^ Growing evidence also shows that the human microbiome is closely related to diverse skin wound healing, including diabetic chronic wounds,^49^ sleep deprivation-induced skin injury ^23^ and UVB-caused skin damage ^24,25^ depending on microbial SCFA production through regulation of the gut-skin axis. However, little information is available regarding the exact mechanisms between the gut microbiome and NM-caused dermal toxicity. In the present study, we identified that BA reduction was the typical characteristic of the gut microbial metabolome in NM-exposure mice. HC-FMT or *A. muciniphila* markedly increased BA levels in the feces and serum, thereby inhibiting NM-induced skin injury. In addition, BA or SCFAs mix also significantly attenuated NM-caused cutaneous damage and the protective effect of the two groups was similar. Latest researches have shown that cyclophosphamide (a widely used NM-derived alkylating agent) decreases SCFA levels resulting in immumosuppression in mice.^59,60^ Our findings clearly indicated that BA was the functional metabolite produced by the gut microbiota, especially *A. muciniphila,* that ameliorated NM-induced skin injury, reshaping our understanding of the pathogenesis and treatment of refractory wounds.

Furthermore, the underlying mechanisms responsible for NM-induced changes of the gut microbiota were further investigated. The gut microbiota consists of diverse microorganisms residing in the gastrointestinal tract and is regulated by various genetic, health, and environmental factors such as α-defensins. α-Defensins are Paneth cell-derived antimicrobial peptides in the intestinal crypts, which require MMP7-mediated proteolytic activation for their bactericidal function.^61^ α-Defensins function as pivotal regulators of the gut microbiota through selective elimination of pathogenic bacteria without harming commensals.^27^ Previous studies have demonstrated that HD5 levels are markedly declined in elderly individuals (>70 years) and correlate negatively with *Alistipes*, but positively with *Lachnospiraceae* (well studied BA-producing bacteria).^62^ Shorter sleep time decreases HD5 secretion, resulting in a decline in SCFA-producing bacteria (e.g., [*Ruminococcus*] *gnavus group* and *Butyricicoccus)* and a concomitant reduction in SCFA generation in humans.^63^ After radiation, the abundance of opportunistic pathogenic bacteria (*Streptococcus* and *Escherichia-Shigella*) is increased at the expense of probiotics (*Lactobacillus* and *Desulfovibrio*), leading to deceased SCFAs in α-defensin deficiency mice.^33^ Neonicotinoid pesticide clothianidin exposure inhibits α-defensin production thereby reducing the abundance of BA-generating bacteria *Lachnoclostridium* and *[Eubacterium] xylanophilum group* in mice.^32^ In our study, we established that NM remodeled the gut microbiota, particularly reducing *A. muciniphila* abundance, leading to a decline in BA contents and subsequent dermal toxicity in an α-defensin-dependent manner. Moreover, it has been demonstrated that VDR is critical for α-defensin secretion. VDR is abundantly expressed in normal gut epithelial cells, displaying a surface-to crypt gradient with maximal levels localized to the crypts.^64^ *Lactobacillus plantarum* increases intestinal VDR expression and Paneth cell numbers, thereby promoting α-defensin secretion, ultimately ameliorating *Salmonella*-caused colitis in vitro and in vivo; while, these protective effects are abolished in the absence of VDR.^37^ Wu et al. cross the flox-*Vdr* mice with p-*villin*-Cre mice to generate specific intestinal *Vdr*-knockout mice and find that α-defensin 5 (DEFA5) expression is significantly decreased in intestinal *Vdr^−/−^* mice, which are more vulnerable to CCL4-caused liver fibrosis.^36^ Another study from Su and his colleagues reported that VDR deficiency inhibits the expression of α-defensin and MMP7, thereby increasing *Helicobacter hepaticus* at the expense of *A. muciniphila;* however, oral administration of DEFA5 notably abolishes VDR deficiency-caused intestinal flora dysbiosis thereby attenuating hepatic steatosis.^35^ Researchers have shown that after dermal exposure, NM is rapidly absorbed and partially retained in the skin, causing localized damage.^65^ The remaining fraction can penetrate into the systemic circulation, leading to systemic toxicity, with particularly severe damage to rapidly proliferating tissues such as intestinal epithelium and bone marrow hematopoietic cells.^2^ In the current study, we identified that dermal exposure of NM reshaped the gut microbiota through inhibiting the intestinal VDR-α-defensin singaling pathway in mice. While the intestinal VDR-α-defensin pathway is known to regulate the gut microbiota in multiple disease models, our study is the first to demonstrate its pivotal role in NM-induced gut microbiota dysbiosis. This work advances understanding of the mechanisms through which NM triggers dysbiosis.

Finally, we demonstrated that the “intestinal VDR-α-defensin-gut microbiota-SCFAs” signaling pathway was involved in VD3’s protective effect against NM-caused skin injury. VD3 is a readily available, fat-soluble secosteroid and serves as a necessary nutrient for maintaining human health by binding to VDR. Accumulating evidence illustrates that VD3 protects against NM-induced cutaneous injury by mitigating inflammation,^39^ inhibiting ferroptosis^40^ and tissue destruction;^66^ however, the underlying mechanisms remain incompletely understood. A recent systematic review of human studies provides strong evidence that VD supplementation markedly changes the gut microbiota composition, with particularly decreased abundance of *Veillonellaceae* and *Oscillospiraceae.*^41^ In a LPS-stimulated systemic inflammation mouse model, VD3 reverses LPS-caused reduction of *Clostridiales_bacterium_CIEAF_020* and inflammation.^43^ VD3 also reshapes the microbiota by enrichment of *Bacteroidetes, Proteobacteria,* and *Parabacteroides* and depletion of *Firmicutes* and *Ruminococcus,* a shift that may contribute to its beneficial effect on obesity in mice.^42^ Zhou et al. clarified that VD3 notably increases *A. muciniphila* abundance, along with a reduction of risk for colorectal cancer in mice.^67^ In the present study, we found that VD3 remodeled the gut microbiome in mice thereby inhibiting NM-caused dermal toxicity, in which *A. muciniphila* seemed to be the core functional bacterium. Meanwhile, targeted metabolomics analysis recognized that VD3 significantly increased BA levels in the feces and serum of NM-treated mice, which was positively correlated with *A. muciniphila* abundance and was abolished by ABX. *A. muciniphila* markedly attenuated ABX-caused reduction of BA levels and BA administration ameliorated ABX-induced inhibition of the VD3’s beneficial effect on NM-caused wounds. These findings suggested that BA could be the potential functional metabolite produced by the gut microbiota especially *A. muciniphila,* and mediated the protective effect of VD3 against NM-induced skin injury. As expected, we also shown that VD3 increased intestinal VDR and α-defensin expression; while, VDR or α-defensin deficiency notably abolished VD3-mediated modulation of the gut microbiota and BA production in NM-treated mice. It has been found that VD3 protects against live injury^35,36^ and colitis,^37^ and alters cancer immunity^44^ via remodeling the gut microbiota through the intestinal VDR-α-defensin signaling pathway. Recently, Liao et al. confirmed that lack of VD3 decreases antimicrobial peptides expression, thereby reducing *Cetobacterium* abundance and acetate levels, ultimately increasing the susceptibility to bacterial infection in zebrafish.^68^ Our present data clearly indicated that VD3 increased *A. muciniphila* abundance and BA contents via activation of the intestinal VDR-α-defensin signaling pathway, suggesting that VD3 was a feasible treatment option for NM-induced dermal toxicity by targeting the gut microbiota.

## 5. Conclusion

To the best of our knowledge, we have, for the first time, uncovered a novel mechanism whereby NM-induced dysbiosis of the gut flora, characterized by a decrease in *A. muciniphila* abundance, promoted dermal injury by suppressing microbial BA production through inhibition of the intestinal VDR-α-defensin signaling pathway. Interestingly, VD3 seemed to exert beneficial effects against NM-caused dermal toxicity by altering the gut flora, particularly increasing *A. muciniphila* content, leading to increased microbiota-generated BA, via activation of the intestinal VDR-α-defensin axis **(Fig. 8)**. Our findings reveal a novel mechanism of VD3-mediated protection against NM-induced cutaneous injury, indicating that targeting the gut microbiota or supplementing with BA could be effective therapies for vesicant-induced skin wounds.

**Figure 8.**
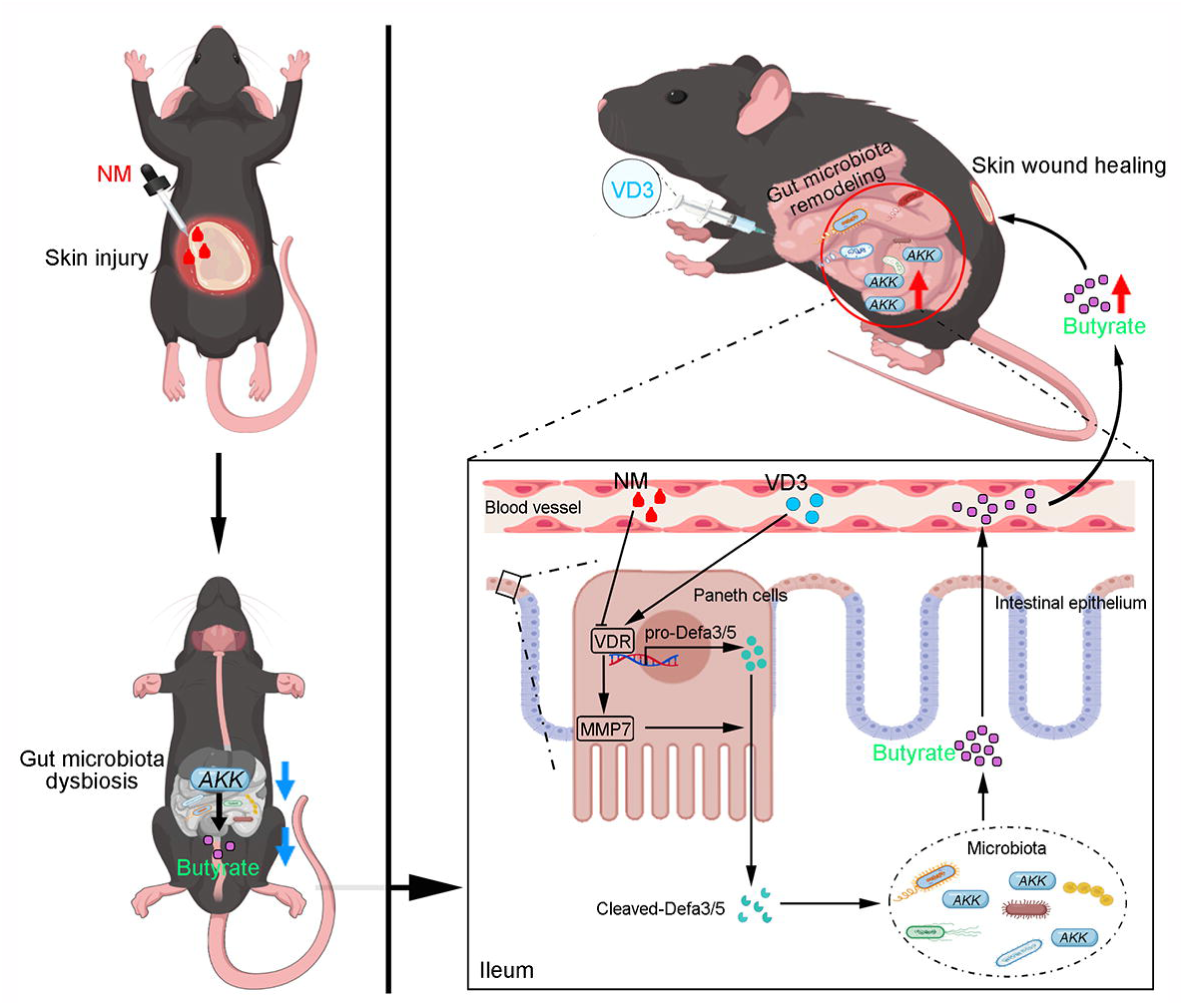
VD3 attenuates NM-induced dermal toxicity by enhancing microbial butyrate production via the intestinal VDR-α-defensin signaling pathway.

## Abbreviations

AAV-*shMmp*: adenovirus-associated virus serotype 9 carrying with a *Mmp7* short hairpin RNA
AAV-*shVdr*: adenovirus-associated virus serotype 9 carrying with a *Vdr* short hairpin RNA
ABX: antibiotics
ACTB: β-actin
*A. muciniphila/AKK*: *Akkermansia muciniphila*
ANOSIM: analysis of similarities
ANOVA: analysis of variance
ATCC: American Type Culture Collection
AUC: area under the curve
BA: butyric acid/butyrate
CFUs: colony-forming units
DAPI: 4’,6-diamidino-2-phenylindole
DEFA5: α-defensin 5
DEGs: differentially expressed genes
*E. coli*: *Escherichia coli*
*E. faecalis*: *Enterococcus faecalis*
FMT: fecal microbiota transplantation
HC-FMT: fecal samples from the controls transplantation
HD5: human alpha defensin-5
LDA: linear discriminant analysis
LEfSe: LDA effect size
LPS: Lipopolysaccharide
MMP7: matrix metalloproteinase 7
NM: nitrogen mustard
OPLS-DA: orthogonal partial least squares discrimination analysis
PCoA: principal coordinate analysis
PBS: phosphate-buffered saline
PLS-DA: partial least squares discrimination analysis
PERMANOVA: permutational multivariate analysis of variance
qPCR: quantitative polymerase chain reaction
ROC: receiver operating characteristic
SCFAs: short chain fatty acids
SD: standard deviation
SI: small intestine
SVM: support vector machines
UV: ultraviolet
VD3: vitamin D3
VDR: vitamin D receptor
VIP: variable importance in projection.

## Data availability

All data and materials are available to the researchers once published.

## Acknowledgments

We are particularly grateful to Figdraw (https://www.figdraw.com/static/index.html) for providing a useful tool to draw the diagram of the mechanism.

## Author Contributions

M. -L.C., Y.H., X.-F.H, X.-H.D, A.-P.W, H.-B.S, and Z.-M.Z were involved in the study design. Y.H., X.-F.H., Z. Z, F.Y, X.-H.D., Z.-Y.P., X.-G.W., J.-Q.Z., G.-R.D, H.T, H.-Y.C. and M.Q. conducted the experiments and the statistical analyses. M.-L.C., Y. H., X.-F.H., X.-H.D, and Z.-M.Z. wrote the first draft of the manuscript. All authors contributed to the final version of the manuscript. A.-P.W, H.-B.S, Z.-M.Z. and M.-L.C. had the primary responsibility of the final content. All authors have read and approved the final manuscript.

**Generative AI and AI-assisted technologies were NOT used in the preparation of this work.**

