## Supplementary Materials for "Vitamin D3 attenuates nitrogen mustard-induced dermal toxicity by enhancing microbial butyrate production via the intestinal VDR-α-defensin signaling pathway"

#Correspondence

**Running title:** VD3 attenuates NM-caused skin injury

**1. Materials and methods**

**1.1 Reagents and antibodies**

Mechlorethamine hydrochloride (NM, K900001X) was purchased from Dibo (Shanghai, China). Acetate (S7545), propionate (P5436) and BA (303410/B5887) were obtained from Sigma-Aldrich (St. Louis, MO, USA). Ampicillin sodium (HY-B0522A), vancomycin hydrochloride (HY-17362), neomycin sulfate (HY-B0470), metronidazole (HY-B0318), and VD3 (HY-15398) were supplied by MedChemExpress (Monmouth Junction, NJ, USA). Antibodies were sourced as follows: anti-MMP7 (10374-2-AP), anti-VDR (10350-AP) and anti-β-actin (ACTB, 66009-1-Ig) were from Proteintech (Wuhan, China).

**1.2 Intestinal gene silencing via superior mesenteric artery-mediated AAV delivery**

To knock down intestinal *Mmp7* and *Vdr* genes in mice, AAV9 vectors were constructed. The AAV vectors were prepared by synthesizing and cloning shRNA sequences targeting mouse *Mmp7* (5′‑ACTCAGAAGACTTCAGTCTTA‑3′) or *Vdr* (5′‑GCTTAAGTCAAGTGCCATTGA‑3′) into the GV687 (pGC-FU-3FLAG-CBh-gcGFP-IRES-puromycin) or GV478 (pAAV-U6-shRNA-CAG-EGFP) backbone vectors (Shanghai Genechem Co., Ltd.), respectively). Recombinant vectors were verified by DNA sequencing. Commercial scrambled shRNA sequences CON539 (TTCTCCGAACGTGTCACGT) and CON305 (CGCTGAGTACTTCGAAATGTC) were used as negative controls for *Mmp7* and *Vdr*, respectively. Viruses were packaged by co-transfecting 293T cells with the viral vectors and helper plasmids pHelper and pRepCap, harvested 72 hours post-transfection, and purified and concentrated via iodixanol gradient ultracentrifugation. Viral titers were determined by qPCR. All AAV viruses were prepared in 0.001% Pluronic F-68 solution, with a final titer of 1×10¹² vg/mL.

Mice were anesthetized with sodium pentobarbital. A midline laparotomy was performed to expose the superior mesenteric artery as previously reported.^1^ After proximal clamping, 100 μL of AAV9 suspension was slowly infused distal to the clamp via a 33-gauge Hamilton syringe over 2-3 minutes. The clamp was maintained for 5 minutes post-injection to limit systemic leakage before reperfusion. The abdomen was closed in layers, and mice were returned to routine housing postoperatively. Mice were monitored daily for body weight and surgical recovery. Four weeks post-injection, mice were treated with VD3 and NM as described above. The intestinal tissues were collected at 7 d after NM-exposure to evaluate the expression of *Defa3*, *Defa5*, MMP7, and VDR by qPCR or western blot analysis.

**1.3 Metagenome** **Analysis**

Fecal samples from mice were collected, snap-frozen in liquid nitrogen, and stored at -80 °C. Metagenomic sequencing was performed on an Illumina NovaSeq 6000 platform (Guangzhou Gidio Biotechnology Co., Ltd., China) using the NEBNext® Ultra™ DNA Library Prep Kit (NEB, USA). Raw reads were filtered with FASTP (v0.18.0) to obtain clean reads. Clean reads from each sample were individually assembled using MEGAHIT (v1.1.2) with k‑mers ranging from 21 to 99. Genes were predicted from the assembled contigs (>500 bp) using MetaGeneMark (v3.38). Predicted genes ≥ 300 bp from all samples were pooled and clustered with CD‑HIT (v4.6) at ≥ 95% identity and 90% coverage to reduce redundancy. Reads were realigned to the predicted genes using Bowtie2 (v2.2.5) to obtain read counts. A final gene catalogue was constructed from non‑redundant genes with > 2 read counts. Taxonomic profiling was performed from clean reads using Kaiju (v1.6.3). Differences in microbial community structure between groups were analyzed by Shannon, Chao1, ACE, and Simpson index, PCoA and ANOSIM test through “vegan” R package. Taxonomic comparisons were analyzed by Welch’s t test or Wilcoxon rank sum test. LDA coupled withLEfSe method was employed to explore the significant different microbes by “microeco” package^2^ between various groups. Gini index, VIP score, and ROC analysis were used to select the most important microbiota. A Meta analysis was conducted to select significant microbiota related to phenotype alteration.^3^ Combined fold change was calculated using average across diverse groups. Features exhibiting consistent directional differences across individual comparisons were included in the meta-analysis.The combined p-value (cp) for cross-group comparisons was derived using Fisher’s method. For each feature type, the following thresholds were applied: p-value < 0.05, cp < 0.05, and FDR < 0.05. The metagenomic data have been submitted to the National Genomic Data Center database under the BioProject accession number PRJCA058790.

**1.4 Targeted Metabolomics** **Analysis**

Fecal samples were analyzed using the Q300 Metabolite Array Kit (Metabo-Profile Biotechnology, Shanghai, China), which targets 310 metabolites across 12 biochemical classes as we previously reported.^4^ Briefly, samples (feces: 5 mg; serum: 20 μL) were mixed with ice-cold methanol containing internal standards, vortexed vigorously, and centrifuged. The supernatant was analyzed by ultra-performance liquid chromatography coupled with tandem mass spectrometry (ACQUITY UPLC‑Xevo TQ‑S, Waters) under the following conditions: C18 column (2.1×100 mm, 1.7 μm), column temperature 40°C, mobile phase A: 0.1% formic acid in water, mobile phase B: acetonitrile/isopropanol, flow rate 0.4 mL/min. Metabolites were identified by comparison with authentic standards (MSI confidence level 1). Data processing, including peak integration and quantification, was performed using QuanMET software (v2.0, Metabo-Profile). OPLS-DA were applied to examine variations in metabolite profiles among different groups, with PERMANOVA used to determine statistical significance. VIP was obtained based on the OPLS-DA model. Differentially expressed metabolites were defined as metabolites with VIP ≥ 1 and P < 0.05, where the choice of univariate test depended on data distribution normality. A Z score represents the standardized distance between an observation and the control group mean, expressed in terms of standard deviations. Different metabolites between groups were calculated by student t test. Three machine learning methods including random forest, SVM , recursive feature elimination of Boruta, and ROC analysis were employed to determine the most significant metabolites.

**1.5 Ileal tissue transcriptomic analysis**

On day 2 post NM exposure, The ileal tissue of mice (a 0.5 cm segment distal to the cecum) in each group were collected and placed in RNAlater solution, and stored at -80°C until analysis. Samples were shipped to Shanghai Applied Protein Technology Co., Ltd. for RNA extraction and transcriptomic sequencing. DEGs were identified using DESeq2 analysis and visualized through a volcano plot. Functional characterization of the DEGs was further performed through GO enrichment analysis, revealing their predominant associations with biological processes, molecular functions, and cellular components. We deposited raw RNA-seq data to the National Genomic Data Center database under the BioProject accession number PRJCA058790.

**1.6 ABX treatment**

A broad-spectrum antibiotic cocktail was administered in drinking water ad libitum starting 7 days before NM exposure. The cocktail consisted of ampicillin (1 mg/mL)， metronidazole (1 mg/mL), neomycin (1 mg/mL), and vancomycin (0.5 mg/mL).^5^ Antibiotic administration did not significantly affect body weight, food intake, or water consumption, as assessed using Promethion metabolic cages.

**1.7 Fecal microbiota transplantation**

Recipient mice were pretreated with or without ABX for 7 days. FMT or bacterial supplementation was performed at Day 1 and Day 4 after NM exposure. Mice were orally gavaged with 200 μL of fecal suspension from donor mice or a live preparation of *AKK* (ATCC BAA835; Mingzhoubio Technology Co., Ltd., China) at 1 × 10⁹ CFU/mL.^5^ Vehicle control mice received an equal volume of phosphate-buffered saline (PBS) on the same schedule.

**1.8 SCFAs intervention**

Following NM exposure, mice were fed a control diet. For intervention, drinking water was supplemented with either a SCFA mixture (67.5 mM acetate, 40 mM butyrate, 25.9 mM propionate) or butyrate alone (100 mM) as described previously.^5^ Treatment continued for 7 days prior to subsequent analyses.

**1.9 Quantification of *AKK* by qPCR**

Fecal samples were collected 48 hours post-NM exposure. Genomic DNA was extracted using using the QIAamp DNA Stool Mini Kit (Qiagen, Venlo, Netherlands) according to the manufacturer's instructions and diluted to 1 ng/µL. Each diluted sample (2 µL) was analyzed in triplicate by qPCR using a 20 µL SYBR Green reaction system (AG11701, Accurate Biology) with primers specific for *AKK* according to the manufacturer’s instructions and was performed on the CFX Connect real-time PCR detection system (Bio-Rad). The quantitation of bacteria was determined relative to the *Actb* gene, as previously reported.^6^ The specific primers used for these groups were indicated in **Table. S1**.

**1.10 SCFAs extraction and measurement**

Fecal and serum samples were stored at −80°C. For SCFA extraction, 30 mg feces were homogenized in 300 μL water, shaken at 4°C for 30 min, and centrifuged. The supernatant (100 μL) or serum (diluted 1:1 with water) was acidified with 5 M HCl to pH ~2 and extracted with diethyl ether. After phase separation and dehydration with anhydrous Na₂SO₄, the ether layer was derivatized with O-bis(trimethylsilyl) trifluoroacetamide and analyzed by gas chromatography–mass spectrometry as described before.^7^ Background correction was performed using parallel-processed blank samples.

**1.11 Western Blot Analysis**

Western blotting was performed as previously described.^8,9^ Ileal tissues were lysed in RIPA buffer containing protease and phosphatase inhibitors. Protein concentration was determined using a BCA assay (Thermo Fisher Scientific). Proteins (60–100 μg) were separated by SDS‑PAGE (10–12% gels) and transferred to PVDF membranes. Membranes were blocked with 5% skimmed milk and incubated overnight at 4°C with primary antibodies against MMP7 (1:1000), VDR (1:1000), and ACTB (1:1000). After washing, membranes were incubated with HRP-conjugated secondary antibodies (Thermo Scientific) and visualized using enhanced chemiluminescence on a Vilber Fusion FX7 system.

**1.12 qPCR analysis of target gene expression**

Total RNA was isolated from ileal tissue specimens using TRIzol™ Reagent (Invitrogen, Carlsbad, CA, USA). Reverse transcription was carried out with the PrimeScript™ RT Reagent Kit (Takara Bio Inc., Kusatsu, Japan), and the resulting cDNA was subjected to qPCR using SYBR Green reaction system (AG11701, Accurate Biology) with primers specific for *Defa3, Defa5,* and *Hd5* according to the manufacturer’s instructions as described above. All primer sequences are provided in **Table. S1**.

**1.13 Immunofluorescent staining**

The mice distal ileum, approximately 1 cm proximal to the cecal end, was carefully dissected and immediately flushed with ice-cold PBS to remove luminal contents, then embedded in optimal cutting temperature compound, snap-frozen in liquid nitrogen, and stored at −80°C. Cryosections (8 μm) were prepared and fixed in 4% paraformaldehyde at room temperature (RT) for 30 minutes, Following fixation, tissues were permeabilized with 0.2% Triton X-100 in PBS for 15 min at RT, and blocked with 5% donkey serum for 1 h. Sections were incubated overnight at 4°C with rabbit anti-MMP7 antibody (1:500). Following PBS washes, Alexa Fluor 488-conjugated secondary antibody (1:500; Invitrogen) was applied for 1 h at RT. Images were captured using a confocal microscope (Zeiss LSM 880) and analyzed with Image J software.

**1.14 H&E staining and histopathological analysis**

At 7 d after NM exposure, skin wounds and nearby tissues were removed and fixed in 4% paraformaldehyde containing 0.1% DEPC for histological analysis. Subsequent preparation for paraffin embedding was performed based on routine protocols. Slice thickness was limited to 5 μm for H&E staining and microscopic evaluation (Carl Zeiss, Germany) of histopathological features, such as epidermal thickness, parakeratosis, epidermal denuding and epidermal death, as described previously.^10^.

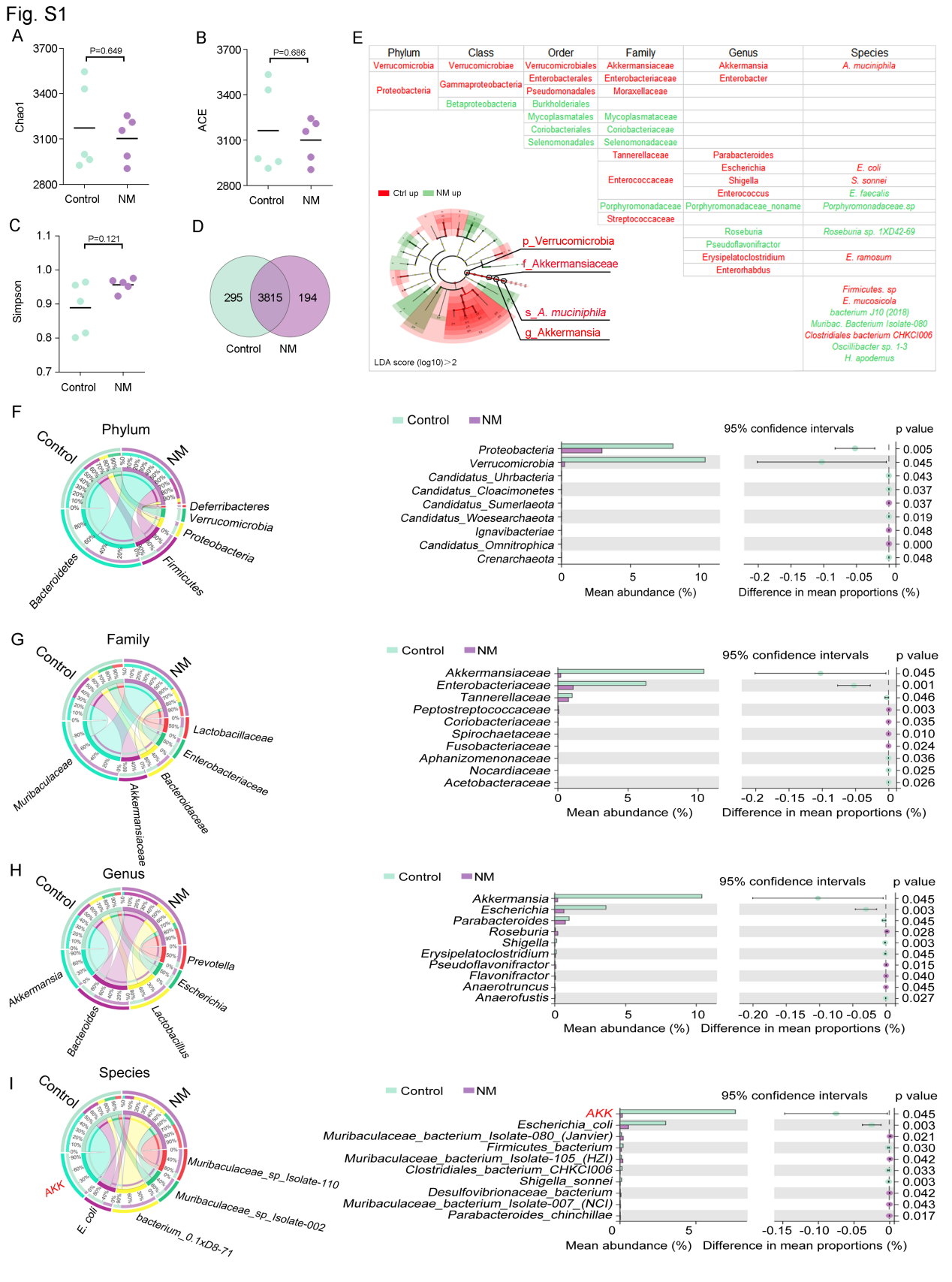

**Figure S1. NM induced dermal toxicity via remodeling the gut microbiota, related to Figure 1.** α-diversity analysis of fecal microbiota in the control and NM groups: **(A)** Chao1, **(B)** ACE, and **(C)** Simpson. **(D)** Venn diagram illustrating shared and unique species between the control and NM group. (**E**) LEfSe analysis of metagenome, taxa enriched in the control (red) and NM (green) groups. Distribution of the predominant dominant gut microbiota at the **(F)** phylum, **(G)** family, **(H)** genus, and **(I)** species levels in Circos (left) and Welch's t-test bar plots (right), respectively. Data are presented as mean ± SD (n = 5 per group).

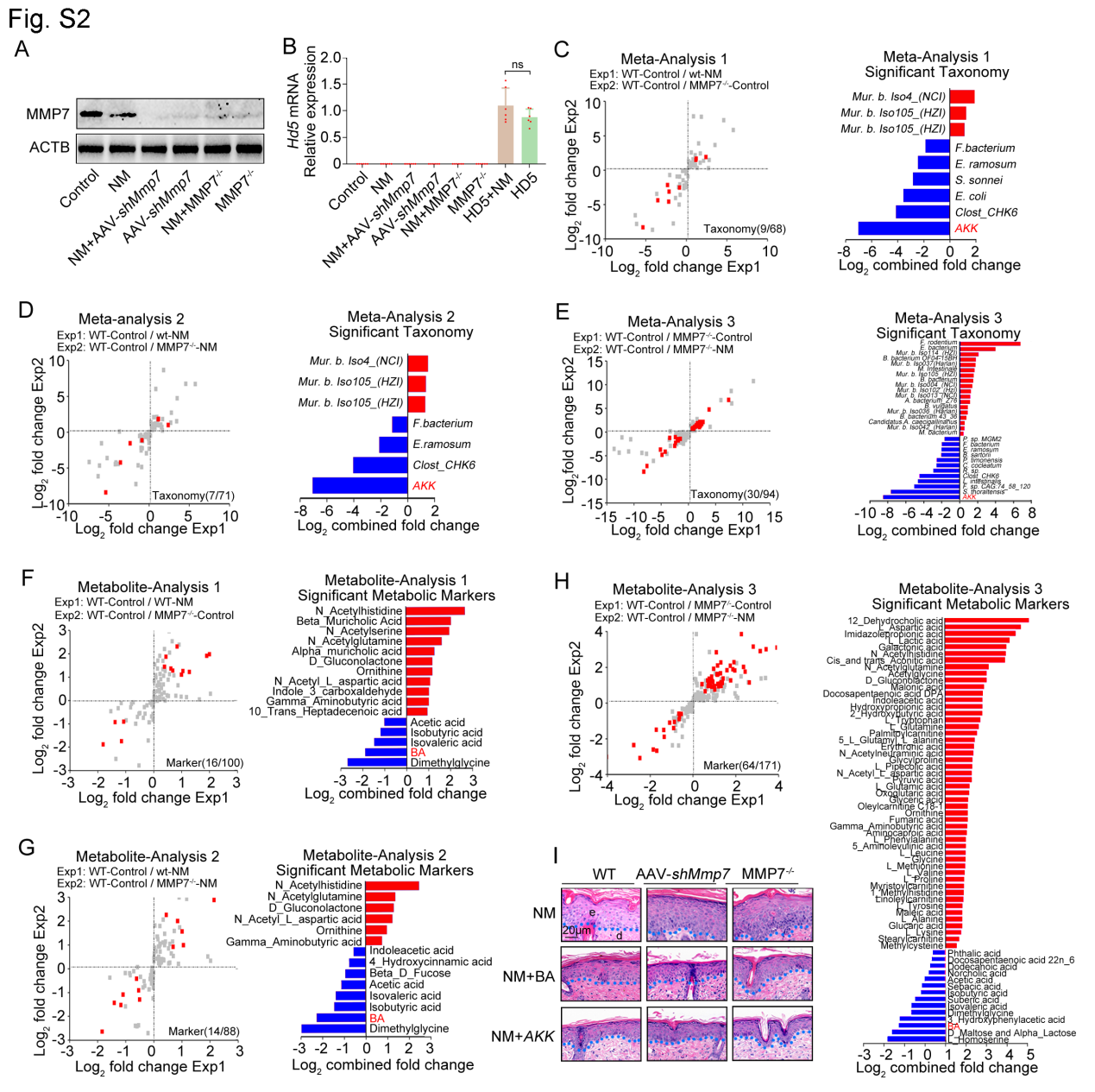

**Figure S2. NM decreased microbial BA levels by inhibiting α-defensin production, related to Figure 3.** (**A**) Western blot analysis of MMP7 expression across the indicated groups (n = 6 per group). (**B**) Relative expression of *Hd5* mRNA in diverse groups (n = 6 per group). (**C-E**) Meta-analysis of metagenomic sequencing data identifying microbial taxa consistently altered in the same direction across datasets (n = 5 per group). (**F-H**) Meta-analysis of targeted metabolomics data revealing metabolites consistently altered in the same direction across datasets (n = 5 per group). (**I**) Representative H&E staining images of skin tissue sections from mice under the indicated conditions (n = 6 per group). Data are presented as mean ± SD.

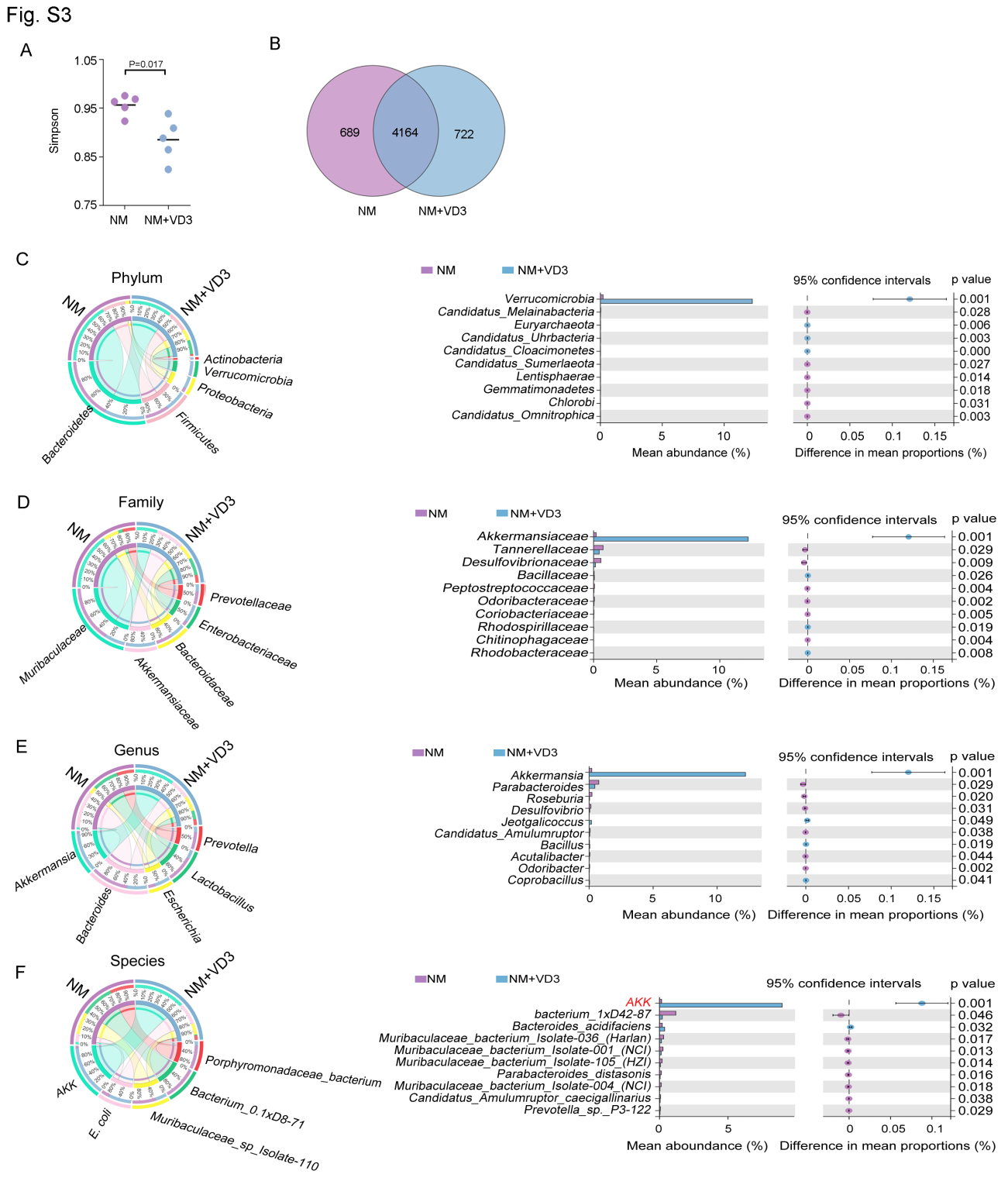

**Figure S3. VD3 attenuated NM-caused dermal toxicity by reshaping the gut microbiota.** **(A)** α-diversity (Simpson index) analysis of fecal microbiota between NM and NM + VD3 group . (**B**) Venn diagram showing the common species between NM and NM + VD3 group. Distribution of the predominant dominant gut microbiota at the **(C)** phylum, **(D)** family, **(E)** genus, and **(F)** species levels in Circos (left) and Welch's t-test bar plots (right), respectively. Data are presented as mean ± SD (n = 5 per group).

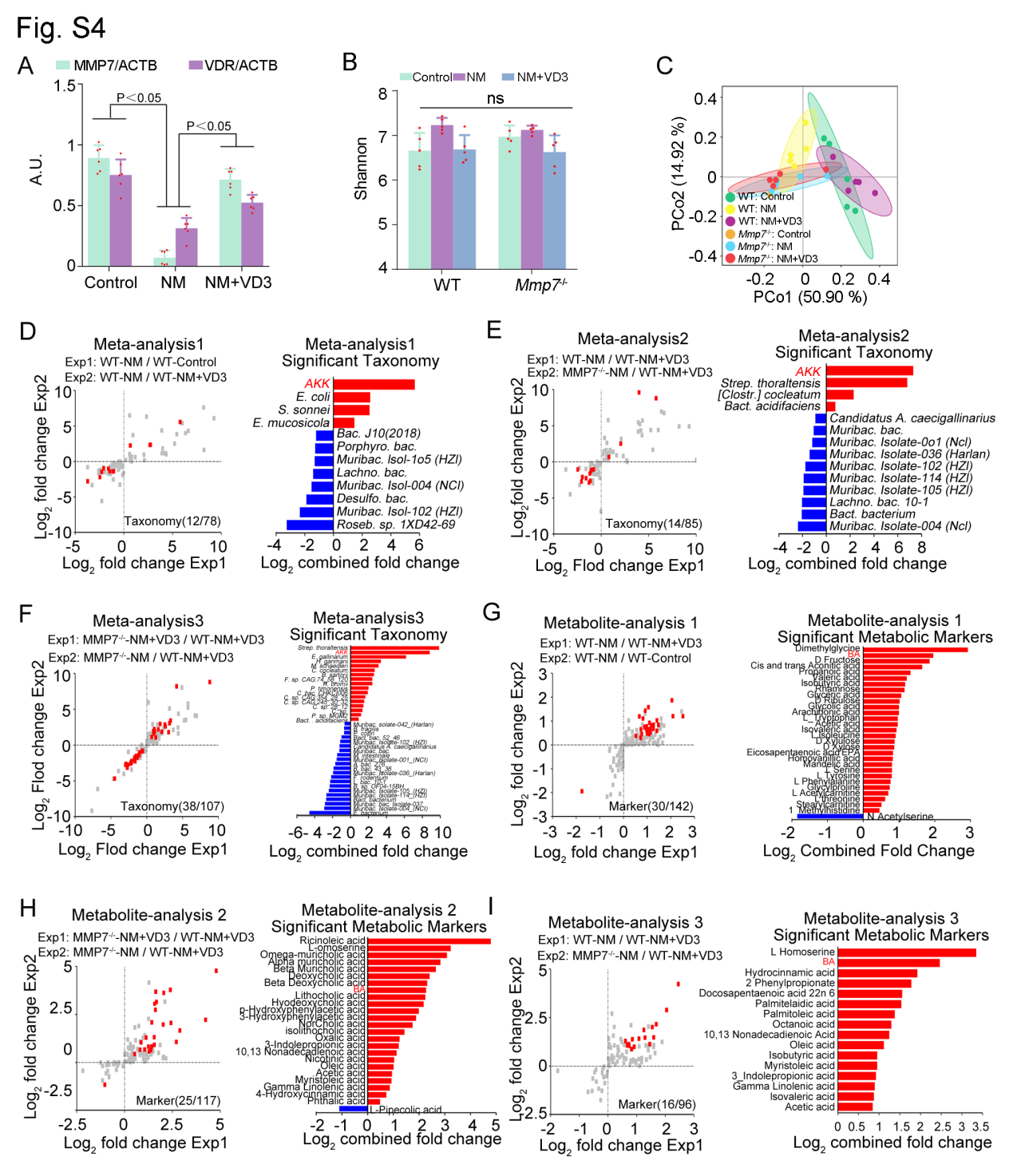

**Figure S4. VD3 increased microbial BA levels via the intestinal VDR-α-defensin signaling pathway, related to Figure 7.** (**A**) Quantification of MMP7 and VDR levels by western blot analysis in the indicated groups, corresponding to **Figure 7B** (n = 6 per group). (**B**) α-diversity (Shannon index) of fecal microbiota at species level in WT and MMP7*^-/-^* mice (n = 5 per group). (**C**) PCoA analysis of metagenome in WT and MMP7*^-/-^* mice with indicated treatments (n = 5 per group). (**D-F**) Meta-analysis of metagenomic datasets identifying microbial taxa consistently altered in the same direction across comparable groups (n = 5 per group). (**G-I**) Meta-analysis of targeted metabolomic datasets revealing metabolites consistently altered in the same direction across the comparable groups (n = 5 per group). Data are presented as mean ± SD.

**Table S1. Primers used for qPCR**

| **Name** | **Primer** | **Sequence (5’ to 3’)** |
| --- | --- | --- |
| *A. muciniphila* | Forward | GACCGGCATGTTCAAGCAGACT |
|  | Reverse | AAGCCGCATTGGGATTATTTGTT |
| *Mouse Defa3* | Forward | TAGTCCTCCTCTCTGCCCTC |
|  | Reverse | ATGACCCTTTCTGCAGGTCC |
| *Mouse Defa5* | Forward | CATTTGTCCTCCTCTCTGCC |
|  | Reverse | GGTCCCAAAAACGCGTTCTC |
| *Human Hd5* | Forward | GCCATCCTTGCTGCCATTC |
|  | Reverse | AGATTTCACACACCCGGGAGA |
| Mouse *Actb* | Forward | GTCCCTCACCCTCCCAAAAG |
|  | Reverse | GCTGCCTCAACACCTCAACCC |
